# Community coalescence altered the potential of horizontal gene transfers within the native soil microbiome

**DOI:** 10.1101/2023.09.18.558195

**Authors:** Xipeng Liu, Thomas Hackl, Joana Falcão Salles

## Abstract

Microbial community coalescence, which refers to the mixing of microbial communities, frequently shapes the assemblage of soil microbiomes in natural ecosystems. It can exert selective pressure on the coalescent taxa, leading to ecological changes in microbial community structure or microbial evolutionary changes via horizontal gene transfer (HGT). However, the influence of community coalescence on the potential of HGTs within native communities, particularly in soil ecosystems, remains poorly understood. Here, we experimentally quantified the potential evolutionary consequences of soil coalescence. We achieved that by subjecting microcosms containing natural soil to invasion by several microbial communities and profiling mobile genetic elements (MGEs) and adaptive genes of microbial communities up to 60 days after coalescence. Our findings revealed both specific and common responses of MGEs to coalescences over time. Specific effects differed across invasive communities and were particularly pronounced in the early stages. Common effects were associated with an increased abundance of insertion sequences (ISs) across different treatments, suggesting that ISs played a crucial role in promoting diversification at the community level. In summary, we showed that changing MGE profiles are an intrinsic response of the soil microbial community to coalescence-imposed pressure. Our study provides new insights into the modulation of adaptability in soil microbial communities by utilizing community coalescences to address global challenges.

## Introduction

Microbial community coalescence, which refers to merging different microbial communities, is a fundamental process on our planet (Rillig et al., 2015). It influences the assemblage of microorganisms, leading to the fundamental changes in composition, succession, and functionalities of the resulting coalescent community (Rillig et al., 2016; Diaz-Colunga et al., 2022; Liu and Salles, 2023). This phenomenon carries significant implications for our understanding of microbial ecology and ecosystem functioning, from the exploitation of beneficial microbial consortia for biofertilization and bioremediation to the investigation of how microbiomes respond to disturbances caused by biotic factors in the context of global changes (Ramoneda et al., 2021; Liu et al., 2023). Therefore, revealing the mechanisms and consequences of community coalescences from various perspectives can provide valuable insights into this biotic disturbance.

Microbial communities respond rapidly to disturbances through ecological and evolutionary processes (Chase et al., 2021; Arnold et al., 2022). From an ecological perspective, previous research has shown that coalescence primarily exerts competitive pressure for resources, which drives compositional and functional changes in the resident community (Liu and Salles, 2023). Specifically, while invasive communities face significant selective pressure, resulting in low survival rates for the invaders (less than 1%), this process can still suppress certain native species, leading to changes in community structure and function (Liu and Salles, 2023). One subsequent consequence is that the surviving taxa benefit from coalescence, exhibiting increased abundance compared to the uninvaded control (Liu and Salles, 2023). However, the evolutionary responses of resident communities to coalescence that poses biotic pressure remain largely unexplored.

Living microorganisms can acquire adaptive traits in changing environments through many ways, such as mutation, gene expression regulation, horizontal gene transfer (HGT), and epigenetic and phenotypic changes (Douglas and Langille, 2019; Pimpinelli and Piacentini, 2020). Among these processes, HGT is a primary mechanism through which microorganisms exchange genetic information with the external environment, connecting them to other individuals. Through the utilization of mobile genetic elements (MGEs) such as plasmids, insertion sequences (ISs), and integrative and conjugative elements (ICEs), HGT facilitates the dissemination of advantageous traits, including antibiotic resistance and metabolic capabilities, which enhance the survival and competitiveness of microorganisms within their respective ecosystems (Woods et al., 2020; Arnold et al., 2022; Horne et al., 2023). Empirical evidence has shown that the rapid evolution of microbes mediated by HGT can be stimulated by abiotic stresses such as antibiotics, pesticides, and irradiation (Pasternak et al., 2010; Hawkins et al., 2019; Pärnänen et al., 2019; Yao et al., 2022). Community coalescence events will likely intensify the selective pressure (Huet et al., 2023; Liu and Salles, 2023), fostering increased interactions and genetic exchange among diverse microbial populations. As a result, these events provide opportunities for transferring genes and genetic elements.

In this study, we aimed to capture indications of microbial gene transfer at a community level during community coalescence. Monitoring evolutionary events and rates in microbes in situ is still challenging at the community level and in the complex context of soils (Brito, 2021; Arnold et al., 2022). We thus primarily employed shotgun metagenomic sequencing and focused on assessing the relative abundance of MGEs in soil, which serves as a proxy for characterizing the potential for horizontal gene transfer (HGT) within communities. Here, we asked the following fundamental question: How does community coalescence influence the MGE profile and the adaptive functions of the resident community over time? We hypothesize that community coalescence will increase HGTs, which will be especially relevant when the resident communities are resilient to the biotic disturbance or are not experiencing an invasion meltdown.

## Methods

### The setup of coalescence experiments

The present study is based on a previous coalescence experiment. In brief, we created nine invasive communities differing in composition and diversity by inoculating diluted soil suspensions (A:10^-1^, B:10^-3^, and C:10^-6^) extracted from three soils (E, M, and L, collected from a salt marsh ecosystem showing a significant difference in composition) in sterile soil for 28 days to allow for soil colonization. Then, coalescence experiments were performed by introducing nine invasive communities adjusted to the same bacterial density (∼5% of the resident community) into a natural soil (i.e., the original/resident community). The details of experiments and soil microbial communities were described in the previous study (Liu and Salles, 2023). In this study, we selected six treatments (E-A, M-A, L-A, L-B, L-C, and uninvaded control) at dates 0, 5, and 60 to evaluate how community coalescences influence the functional traits of the soil microbial community.

### Shotgun metagenomic sequencing and bioinformatic processing

Fifty-four total genomic DNA samples were submitted for library construction and shotgun sequencing at BGI TECH SOLUTIONS (HONGKONG) CO., LIMITED, Hong Kong. The Short-Insert library was used and sequenced on a 2 × 150 bp DNBseq platform. Raw reads were preprocessed by removing adaptor sequences, contamination, and low-quality reads with SOAPnuke (Chen et al., 2018) to obtain clean reads (39,615,462 reads per sample on average). Detailed information on the sequence data is summarized in Supplementary Table S1.

Megahit v. 1.2.9 was used to perform *de novo* assembly for each sample with the k-mer length increasing from 21 to 149 in steps of 20 (Li et al., 2015). Assembled contigs over 500 bp were submitted to Prodigal v.2.6 to predict the protein-coding genes (Hyatt et al., 2010). After discarding genes shorter than 100 bp, genes from all 54 samples were clustered at ≥95% identity and ≥90% overlap with MMseqs2 (Steinegger and Söding, 2017), resulting in a catalog containing 36,890,547 non-redundant genes. Paired-end reads of each sample were mapped to the gene catalog using Salmon v.1.9.0 (Patro et al., 2017) to obtain the relative abundance (estimated as Transcripts Per Million, TPM) of each gene, and the relative abundance was used in downstream analyses.

### Identification of mobile genetic elements, antibiotic resistance genes, and CNPS-cycling genes

The non-redundant gene sequences constructed from metagenomic data were used to identify MGEs. The ORFs were aligned against corresponding databases for annotating plasmids (PLSDB, (Galata et al., 2019)), insertion sequences (ISs, ISfinder (Siguier et al., 2006)), prophages (PHASTER, (Arndt et al., 2016)), transposons (VRprofile2, (Wang et al., 2022)), integrons (INTEGRALL (Moura et al., 2009)), integrative and conjugative elements (ICEs, ICEberg 2.0 (Liu et al., 2019)), and integrative and mobilizable elements (IME, ICEberg 2.0 (Liu et al., 2019)), respectively. Moreover, a manually curated database of MGEs (mobileOG-db, (Brown et al., 2022)) was used to obtain high-quality and functional annotations. The MGE genes are categorized into five functional divisions: integration and excision (IE), phage-related processes (Phage), replication/recombination/repair (RRR), stable/transfer/defense (STD), and transfer (T). The filtering parameters for these annotations were identity ≥ 70%, alignment length ≥ 25 bp, and e-value ≤ 10e-10.

To identify potential antibiotic resistance genes (ARGs), the representative sequences were aligned against the Comprehensive Antibiotic Resistance Database (CARD) (Alcock et al., 2020) using Diamond Blastp (Buchfink et al., 2021). For the functional genes related to carbon, nitrogen, phosphorus, and sulfur cycling, we aligned ORFs against the Carbohydrate-Active EnZymes database (CAZy) (Cantarel et al., 2009), NCycDB (Tu et al., 2019), PCycDB (Zeng et al., 2022), SCycDB (Yu et al., 2021). The cut-off of these annotations was identity ≥ 70%, alignment length ≥ 25 bp, and e-value ≤ 10e-10. The composition and abundance of these genes in invasive and original resident communities are shown in Supplementary Figure S1-4.

### Identifying the number of neighbors of intermediate contigs

The number of neighbors of intermediate contigs refers to the count of other contigs that share overlapping regions or have alignment relationships with a particular intermediate contig. Thus, the number of neighbors of intermediate contigs in different sequenced genomes may indicate the comparable complexity caused by repeated or duplicated regions due to HGT when the sequencing depth and assembly error were similar across samples. In this study, we developed an approach based on contig networks to calculate the ratio of neighbors (number of neighbors per contig) for each sample based on an intermediate assembly graph outputted from Megahit v. 1.2.9 (Li et al., 2015) to further analyze the HGT processes. In brief, the FASTG files were first created from intermediate contigs constructed from 41-mers with the core function “contig2fastg” in Megahit v. 1.2.9. The connection information of assembly graphs was then extracted for neighbors counting and ratio calculation.

### Statistical analysis

The total relative abundance of MGEs across treatments was shown as a Z-score, and the difference in total relative abundance related to the uninvaded control was estimated using a t-test combining an ANOVA test for global variance. To reveal the compositional difference of functional traits (e.g., MGEs, ARGs, and CNPS-cycling genes) between different treatments, we performed PERMANOVA using the Adonis test (999 permutations) with R package “Vegan”. The relative abundance difference of each MGE family compared to the uninvaded control was estimated using the R package “edgeR” and *p* < 0.05 as the significant level.

Spearman’s correlation-based network analysis revealed the potential relationship between MGEs and other elements such as ARGs, CNPS-cycling genes, and ASVs on Days 5 and 60. The strict cut-offs (Spearman’s r > 0.7 or < –0.7 and *p* < 0.001) were used for these analyses. Only nodes and edges significantly positively correlated with MGEs were kept in the network. Networks were visualized with Geiphi 0.9 (Bastian et al., 2009). Moreover, we employed the Erdos-Renyi model (function *erdos.renyi.game* in R package “igraph”) to construct two random networks according to the corresponding empirical networks. The difference in degree distribution between random and empirical networks was estimated with the Kolmogorov-Smirnov test.

We assessed the initial functional (nutrient-cycling genes) similarity between invasive and resident communities before coalescences by calculating Bray-Curtis dissimilarity (1-dissimilarity) with the R package “Vegan.” The Bray-Curtis dissimilarity in MGE composition between coalescent community and uninvaded control was used to infer coalescence impact on MGE profile.

## Results

### Effects of coalescence on community functional traits

The functional traits of coalescent communities depended on time and treatments (Figure 1). Time was the main factor influencing the composition of the overall functional trait (Figure 1a), which was consistent with the bacterial community composition over two sampling times (Figure 1b; *p* < 0.001, Procrustes analysis). The treatment (i.e., invasive communities) only significantly influenced the overall functional trait on Day 5 (*p* < 0.01, Adonis), whereas the functional trait on Day 60 showed no significant differences (*p* > 0.05, Adonis). Such phenomenon was also observed for three specific functional traits, including CNPS-cycling genes, MGEs, and ARGs (Supplementary Figure S5).

**Figure 1.**
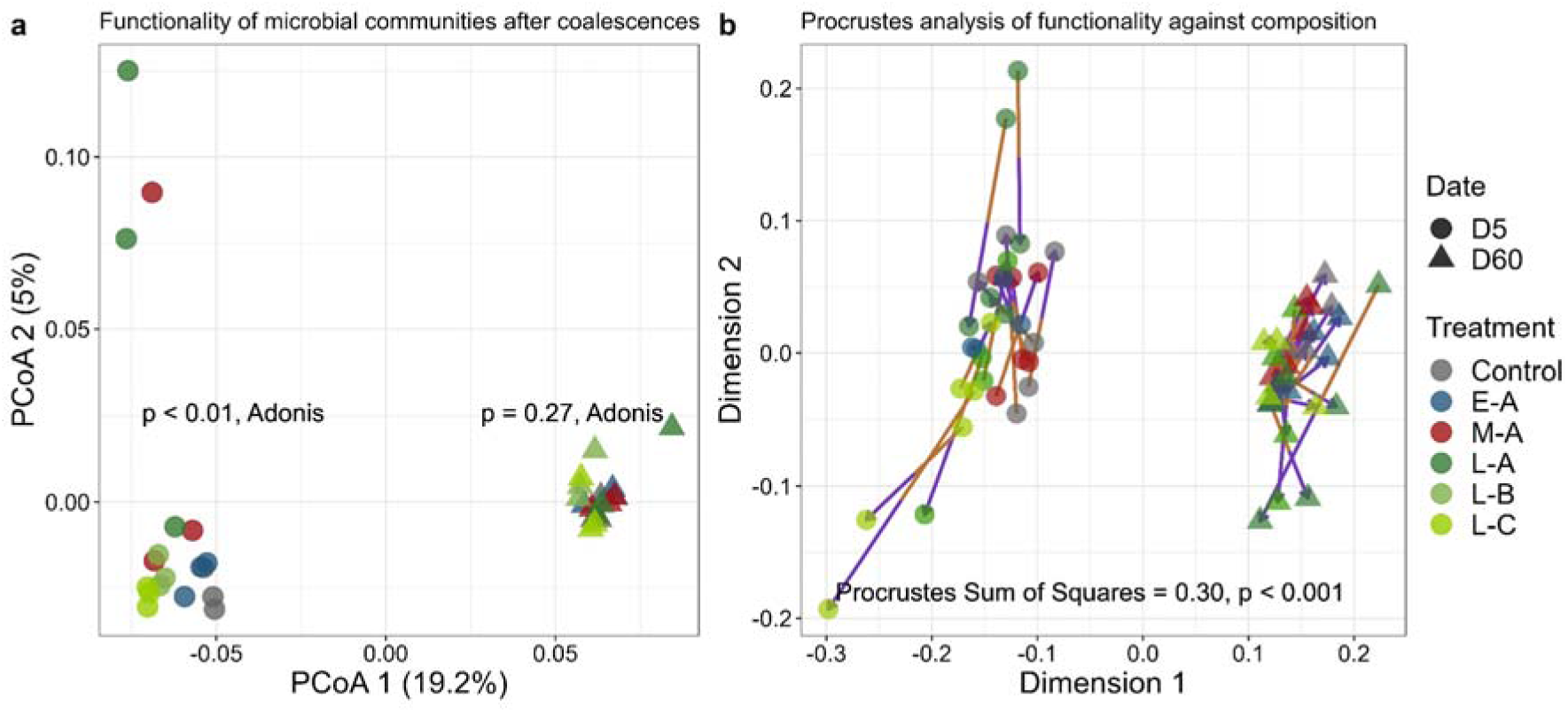
The functionality of microbial communities after coalescence is primarily influenced by time. (**a**) Principal Coordinates Analysis (PCoA) showing the overall functionality (including genes related to nutrient cycling, MGEs, and ARGs) of coalescent communities was mainly changed at the early stage after coalescences. The *p*-value of Permutational Multivariate Analysis of Variance (Adonis) close to different points indicated the functional variance for each date. (**b**) Procrustes analysis of functionality against microbial community composition (based on weighted Unifrac distance) showed that changes in functionality over time were synchronized with community changes. The purple end of each line connects to the 16S rRNA gene data for the sample, while the green end is connected to the overall functionality data.

### Community coalescences altered the profile of MGEs

In the analyzed dataset, mobile genetic elements (MGEs) represented a relative abundance of 4.07-4.81% of all genes (Figure 2). Among the MGEs, plasmids were the most abundant, accounting for an average relative abundance of 3.64% of all coalescent communities (Figure 2). This was followed by ISs at 0.45%, ICEs at 0.29%, prophages at 0.078%, IMEs at 0.020%, transposons at 0.013%, and integrons at 0.0056%.

**Figure 2.**
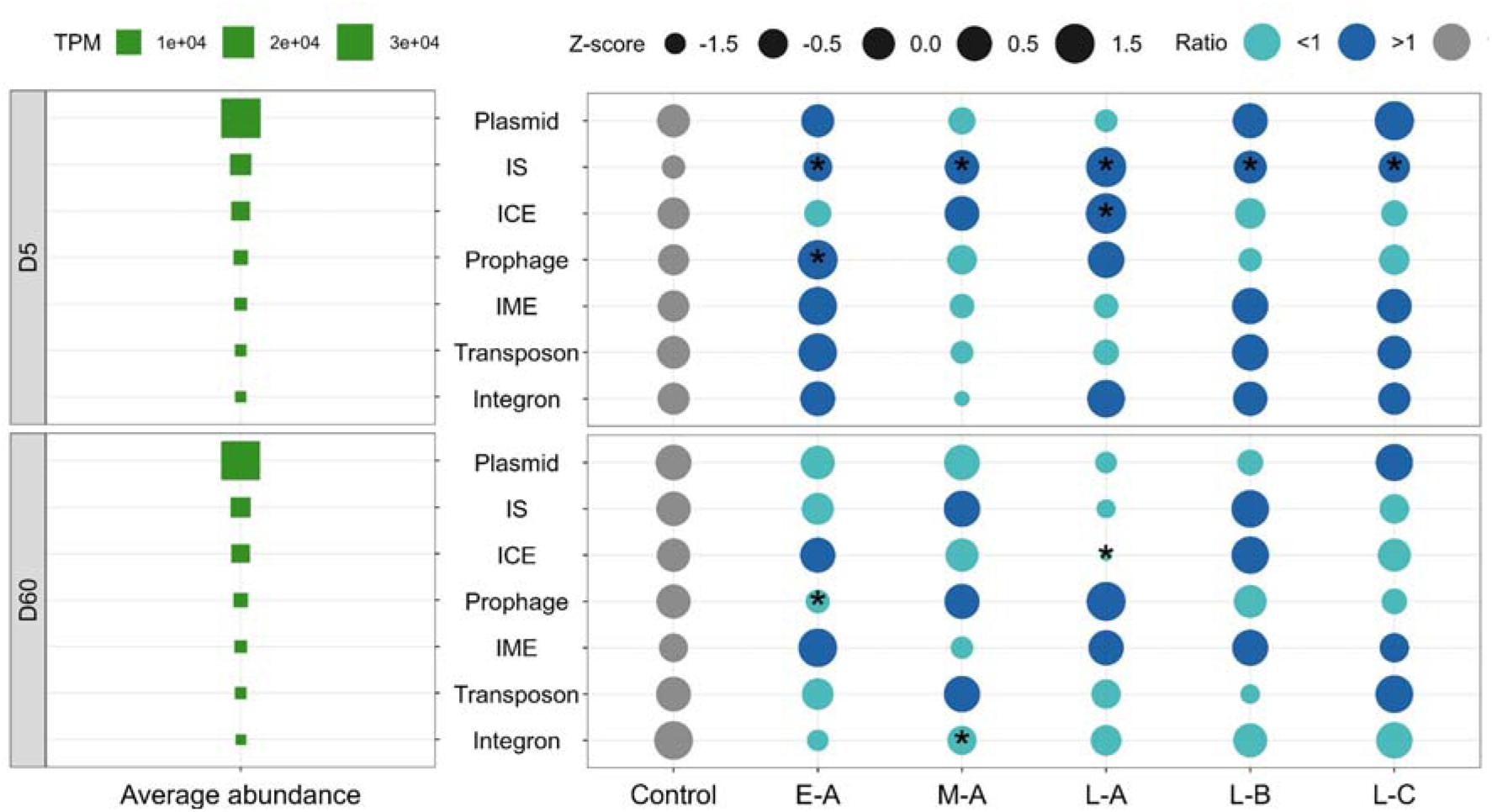
**Coalescences changed the MEG profile of the soil microbial community**. Increased abundance of ISs was the common effect across invasive communities. The abundance (TPM) of MGEs was transformed with Z-score. The color indicates the ratio of MGE abundance in relation to the uninvaded control. Asterisks suggest the significant difference in MGEs’ abundance between uninvaded control and coalescent communities with a t-test (*p* < 0.05).

Different MGEs had varied responses to different invasive communities (Figure 2). On Day 5, the abundance of ISs exhibited the highest sensitivity to coalescence, with all treatments showing a significant increase in IS abundance compared to the uninvaded control on Day 5. The abundance of ICEs was significantly increased in the L-A treatment, and prophage abundance significantly increased in the E-A treatment on Day 5. On Day 60, a significant decrease was observed in several treatments compared to the uninvaded control. For example, ICE abundance decreased significantly in the L-A treatment, prophage abundance reduced considerably in the E-A treatment, and integron abundance decreased significantly in the M-A treatment. However, the abundance of all MGEs in the uninvaded control treatment was not significantly different between D0 and D5 (Supplementary Figure S6). A significant decrease in MGE abundance in the uninvaded control treatment between D0 and D60 was observed for IS and IME (Supplementary Figure S6).

Identifying MGE genes using the MobileOG database allowed for functional classification (Supplementary Figure S7). Among the five critical divisions of MobileOG, the genes encoding proteins for integration and excision (IE) were the most abundant. Furthermore, the total relative abundance of these IE genes showed a significant increase in all coalescent communities compared to the uninvaded control on Day 5. The abundance of the phage-related process genes (P) was significantly increased in the E-A treatment but decreased in the M-A treatment. Other divisions, such as replication/recombination/repair (RRR), stable/transfer/defense (STD), and Transfer (T), were not significantly affected by coalescence on Day 5. For Day 60, the abundance of phage-related process genes decreased in both L-A and L-B treatments. Gene abundance mediating element transfer significantly increased in E-A and L-C treatments. In contrast, the abundance of the STD gene was only observed to increase in the M-A treatment (Supplementary Figure S7). The abundance of all functional groups of MGEs in the uninvaded control treatment was not significantly different between D0 and D5. In contrast, a significant decrease in abundance was observed for all functional groups between D0 and D60 (Supplementary Figure S8).

We further performed correlation analyses for the total abundance of MGEs in invasive communities and the corresponding coalescent communities to assess whether the invader-derived MGEs contributed to the abundance changes after invasions. The results showed the non-significant or negative correlations between the abundance of seven MGEs in invasive communities and coalescent communities for both Days 5 and 60, except plasmid on Day 5 (Supplementary Figure S9). This indicate that the increases in MGEs abundance in coalescent communities, especially for ISs (negative correlation, *p* = 0.0046, Spearman’s correlation, Supplementary Figure S9), were not mainly due to the introduction of invaders.

The total abundance of the integration and excision (IE) division on Day 5 was significant and negatively correlated with its abundance in invasive communities (*p* = 0.048, Spearman’s correlation, Supplementary Figure S10), while the correlation for Day 60 was non-significant (*p* = 0.61). Besides, a positive and significant correlation was found for the stable/transfer/defense (STD) division on Day 5 (*p* = 0.042; the correlation for Day 60 is non-significant with a p-value = 0.046).

### Initial functional similarity between invasive and resident communities predicted MGEs’ response

The impact of coalescence impacts on MGE profiles estimated with the Bray-Curtis dissimilarities between the coalescent community and uninvaded control on Day 5 was positively correlated to the initial functional similarities between invasive and resident communities (Figure 3a). Specifically, among the MGEs, the highest correlation coefficient was observed for IS (R = 0.63) and prophage (R = 0.65). From a functional taxonomic perspective, the genes’ similarity in the carbon and nitrogen cycles exhibited stronger predictability of the post-coalescence effects on MGE profiles. However, such correlations on Day 60 shifted to predominantly negative or non-significant relationships (Figure 3a).

**Figure 3.**
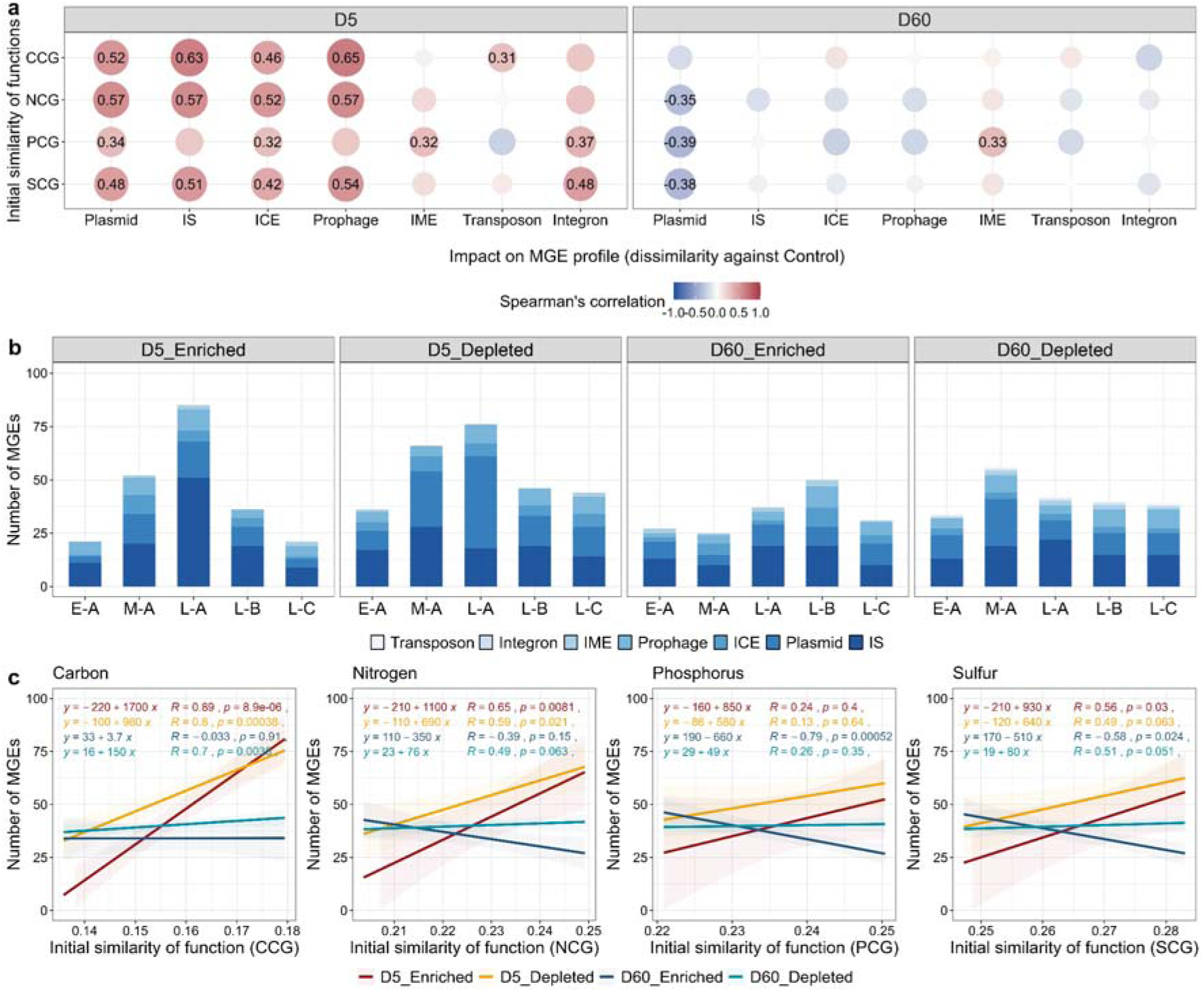
Initial functional similarity between invasive and resident communities correlated with the impacts on MGE profiles suggesting the importance of invader-resident competition. (**a**) Spearman’s correlations between the initial functional similarities between invasive and original resident communities and the impacts on MGE profiles after coalescences. Correlation coefficients (R value) are shown above points, whereas the point without coefficient indicates a non-significant correlation. (**b**) The number of MGEs with significantly changed abundance compared to the uninvaded control on Days 5 and 60. (**c**) Spearman’s correlations between the initial functional similarities between invasive and original resident communities and the number of MGEs with significantly changed abundance. Four functional traits include genes involved in carbon (CCG), nitrogen (NCG), phosphorus (PCG), and sulfur (SCG) cycling.

We obtained the MGEs at the gene family level with significantly enriched or depleted abundance compared to the uninvaded control (Figure 3b and Supplementary Figures S11a and S11b). First, our results revealed that enriched MGEs under different treatments exhibited higher uniqueness (i.e., fewer proportions shared among multiple treatments) for both Days 5 and 60, compared to the depleted MGEs (Supplementary Figure S11c). Besides, we found that IS was the most sensitive MGE type, and the number of increased IS families was higher than that of other MGE types, such as plasmid and ICE (Figure 3b).

The correlation analysis suggested that the total number of enriched MGEs on Day 5 was positively correlated to the initial functional similarities between invasive and resident communities before coalescence (R = 0.89 and *p* < 0.001 based on C-cycling function similarity, R = 0.65 and *p* = 0.0081 based on N-cycling function similarity, R = 0.24 and *p* = 0.4 based on P-cycling function similarity, R = 0.56 and *p* = 0.03 based on S-cycling function similarity, Figure 3c). Such positive correlations were also observed for the total number of depleted MGEs on Day 5 (R = 0.80 and *p* < 0.001 based on C-cycling function similarity, R = 0.59 and *p* = 0.021 based on N-cycling function similarity, R = 0.13 and *p* = 0.64 based on P-cycling function similarity, R = 0.49 and *p* = 0.063 based on S-cycling function similarity). However, the correlation between the total number of enriched/depleted MGEs on Day 60 with initial functional similarities was mostly insignificant or negatively significant (Figure 3c).

### ISs contributed to the number of neighbors of intermediate contigs

We created the intermediate assembly graph (Kmer = 41) for each community and used the ratio of neighbors per intermediate contig (ratio or ratio value hereafter) to infer the potential HGT level in a community, with larger the ratio values indicating more HGT occurred. The reason for using a Kmer size of 41 is that the sequence information can be largely preserved, and the contigs can be obtained in relatively longer lengths (Supplementary Figure S12). The results showed that the uninvaded control had a similar ratio of neighbors (0.23 on average) between Days 5 and 60, indicating the constant status of the HGT event (Figure 4a). However, the coalescent communities had a significantly higher ratio than the uninvaded control on Day 5. M-A, L-A, L-B, and L-C had a similar ratio value and higher than that under the E-A treatment (Figure 4a). On day 60, the ratios for all treatments returned to levels similar to the uninvaded control (Figure 4a).

**Figure 4.**
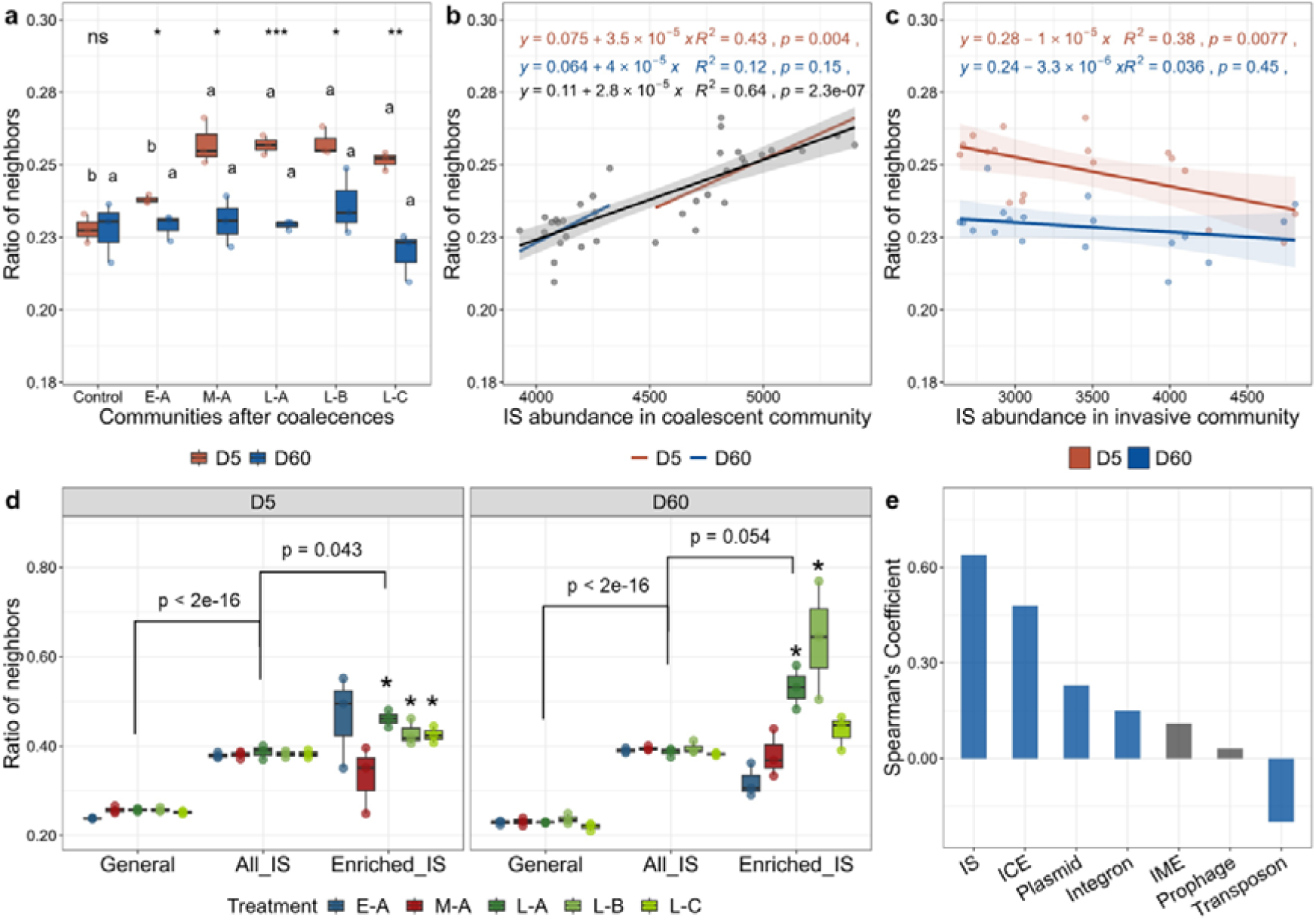
ISs contributed to the increased potential of horizontal gene transfer during coalescences. (**a**) The general ratio of neighbors of contigs in the intermediate assembly graph. Asterisks indicate the significant difference in the ratio between Days 5 and 60, while different letters above the boxes indicate the significant difference among treatments. (**b**) Spearman’s correlation between the ratio of neighbors and IS abundance in coalescent communities. (**c**) Spearman’s correlation between the ratio of neighbors and IS abundance in invasive communities. (**d**) The ratio of neighbors for three contig types: all contigs (General), contigs containing ISs (all_IS), and contigs containing enriched IS families (Enriched_IS). P-values (paired t-test) imply the difference in average ratio between groups, and the asterisks above boxes indicate the significantly higher proportion in the group (Enriched_IS) compared to the other two groups (ANOVA with Tukey test, *p* < 0.05). (**e**) Spearman’s correlation coefficients of general ratio value and the abundance of each MGE type. The positive and negative coefficients imply positive and negative correlations, respectively. The grey color of the bar indicates a non-significant correlation (*p* > 0.05).

The total abundance of ISs in coalescent communities was significantly and positively correlated to the ratio value (R^2^ = 0.64, *p* < 0.0001, Figure 4b), which indicated the importance of ISs in response to the coalescence. Importantly, we found that such a ratio value was negatively correlated to the IS abundance in the corresponding invasive communities on Day 5 (R^2^ = 0.38, *p* = 0.0077) and not significantly correlated to that on Day 60 (*p* = 0.45, Figure 4c). This suggests that the increase in ratio values in coalescent communities was not due to the introduction of ISs derived from invaders.

We further compared the ratio value across all contigs (General), contigs containing ISs (All_IS), and contigs containing enriched IS families (Enriched_IS). We found that contigs containing enriched ISs had the highest average ratio value (Figure 4d). Specifically, the average ratio value of the Enriched_IS group was significantly higher than that of the other two groups on Day 5 (Figure 4d; *p* < 0.05, paired t-test). For Day 60, the average ratio value of all treatments was assessed with a non-significant difference between enriched_IS and all_IS groups with a *p*-value = 0.054 (paired t-test). However, the ratio values of enriched_IS contigs under L-A and L-B treatments were significantly higher than that of the other two groups (*p* < 0.05, ANOVA with Tukey test) for both Days 5 and 60 (Figure 4d).

The overall contribution of MGEs to the general ratio of contigs’ neighbors was estimated with Spearman’s correlation between the ratio value and MGE abundance (Figure 4e). The result showed that ISs had the highest coefficient (R^2^ = 0.64) with the ratio value, followed by ICE (R^2^ = 0.48), plasmid (R^2^ = 0.23), and integron (R^2^ = 0.15). The correlations were non-significant for IME (R^2^ = 0.11) and prophage (R^2^ = 0.03) and negative for Transposon (R^2^ = 0.20).

### Network analysis revealing potential associations to MGEs

Two correlation-based networks revealed the difference in the potential linkages between MGEs and ASVs, ARGs, and CNPS-cycling genes on Days 5 (Network D5) and 60 (Network D60) (Figure 5a). The degree distribution of these two networks was estimated to significantly differ from the corresponding random networks (*p* < 0.001, Kolmogorov-Smirnov test, Supplementary Figure S13). For networks D5 and D60, the total number of positive links was 5320 and 4726, the average degree was 1.29 and 1.18, the average path length was 10.38 and 12.59, and the maximum degree of the node was 34 and 17, respectively (Figure 5b).

**Figure 5.**
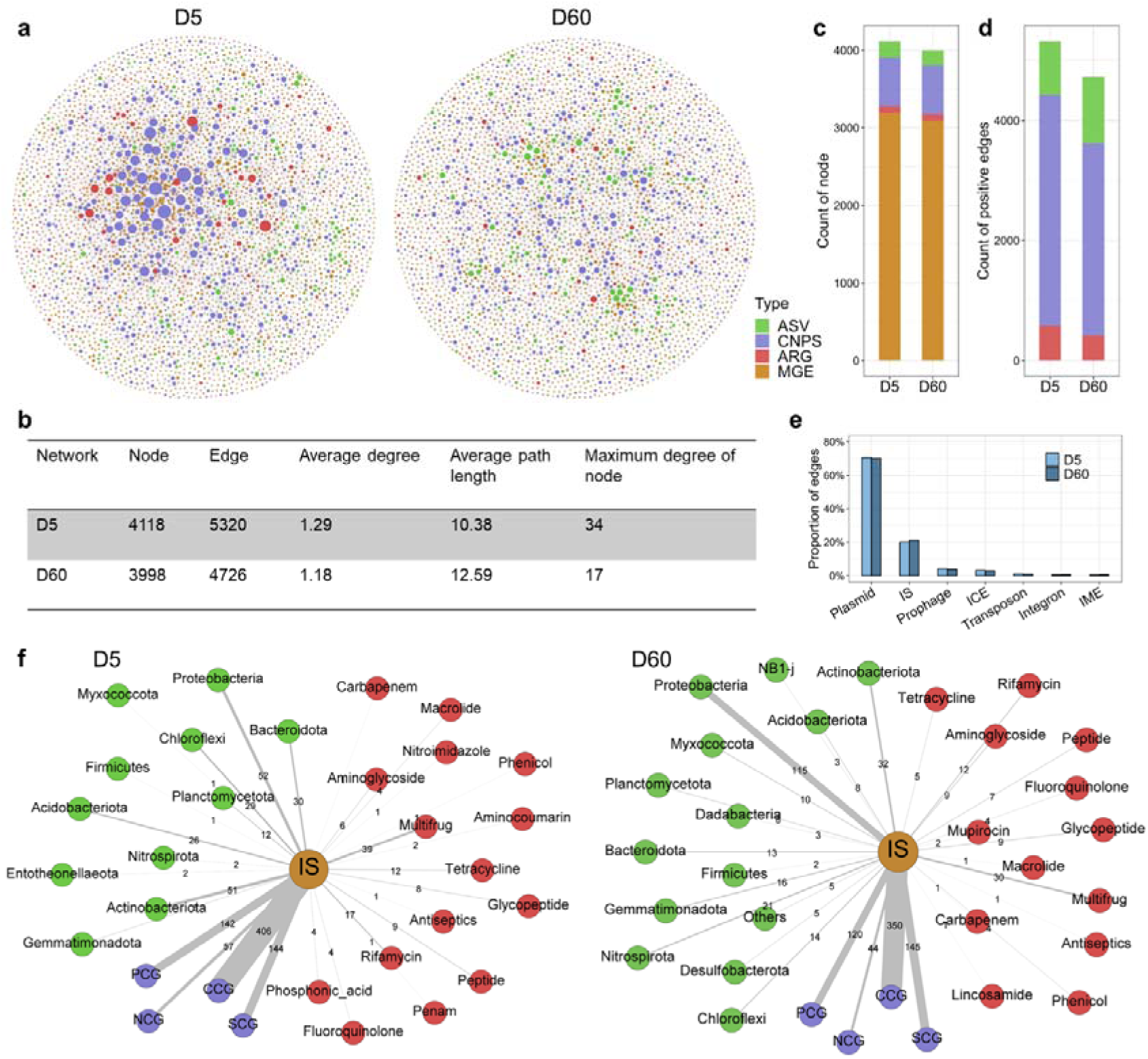
Correlation-based network analyses on the potential associations to MGEs. (**a**) Networks (for Days 5 and 60) based on four types of components: MGEs, nutrient-cycling genes (carbon, nitrogen, phosphorus, sulfur, CNPS), ARGs, and microbial taxa (ASVs). Only significant and positive correlations (Spearman’s r > 0.7 and *p* < 0.001) are shown in the network. The nodes’ size represents the degree number and is scaled to the maximum number of degrees. (**b**) Network metrics. (**c**) The total number of four components in two networks. (**d**) The total number of edges of ARGs, CNPS, and ASVs linked to MGEs. (**e**) The proportion of edges between MGEs and other components in a total number of edges. (**f**) Sub-networks show significant and positive associations with ISs. The width of the line represents the association amount and is indicated by the number next to the line.

The node distribution of four components was similar between the two networks, although the total number of nodes was higher in the D5 than in the D60 network (Figure 5c). The MGE-ARG and MGE-CNPS links were 575 and 3850 on Day 5, respectively (Figure 5d). These were higher than those on Day 60 (411 and 3209, respectively) under the same network construction approach. For example, the total positive correlations between MGEs and the multidrug group of ARGs accounted for 201 (3.8% of total positive edges) and 118 (2.5% of total positive edges) on Days 5 and 60, respectively. However, the total number of positive links between MGEs and ASVs was observed with a different phenomenon, while such links for network D5 (895) were lower compared to network D60 (1106) (Figure 5d).

Plasmids had the highest positive correlations with other components, while such proportion slightly decreased across time (70.6% in D5 and 70.1% in D60, Figure 5e). The proportion of positive edges linked with ISs accounted for the second, which was 20.1% in D5 and 21.2% in D60. By comparing the ratio of positive edges in networks D5 and D60, we found that the proportion increase in D60 was observed for ISs, and this phenomenon did not occur in other major components, such as plasmid, prophage, and ICE. This result may indicate that the importance of IS-mediated associations increased over time after coalescence.

The sub-networks centered with ISs were constructed and showed distinct patterns between D5 and D60 (Figure 5f). Our results suggested that more ARGs and CNPS-cycling genes were correlated with ISs in network D5. For instance, 15 types of ARGs (regarding drug resistance) were found in sub-network D5, compared to sub-network D60 (13 types). Moreover, CNPS-IS edges were 13.7% higher in sub-network D5 (749) than in sub-network D60 (659). However, 11 phyla of microbes (212 in total) correlated with ISs in sub-network D5, which was higher than that in sub-network D60 (14 phyla, 255 total). For instance, taxa affiliated with Proteobacteria were the most frequently present in networks where the number of edges was 52 and 115 in sub-networks D5 and D60, respectively.

## Discussion

Microbial communities are sensitive and adaptive to environmental changes. Due to their fast reproduction, such ecological and evolutionary responses could happen quickly, even within several hours or days (Jain et al., 2003; Bosshard et al., 2017; Dößelmann et al., 2017). The previous attention had been put mainly on either ecological changes of microbial communities or the evolution of specific individuals under abiotic selections. However, the assessment of the microbial evolutionary response under the community coalescence, which represents a common biotic pressure in the soil, is lacking. This study provides vital insight in this respect.

### Specific and common impacts of coalescences on MGEs

Horizontal gene transfer (HGT) is one of the most critical processes in microbial evolution that allows a remarkable increase in genome innovation that significantly exceeds anything that could have been accomplished by clonal evolution alone (Jain et al., 2003). By assessing the relative abundance and composition of mobile genetic elements (MGEs), we demonstrated that coalescences led to alterations in MGE profiles, with the most pronounced changes observed on Day 5 after coalescence. In contrast, the profiles on Day 60 showed resilience compared to the uninvaded control. These findings provide evidence of coalescence-induced modifications in HGT potentials within the soil community.

The influence of coalescence on MGE profiles varied among different treatments. For instance, treatment E-A showed an increase in prophage abundance, while treatments L-A exhibited an increased total abundance of ICE (Figure 2 and Supplementary Figure S9), indicating that the mechanisms for coalescence regulating MGEs are complicated and dependent on invasive communities. Notably, we introduced invaders at a low invasion rate, approximately 5% of the resident taxa density. This suggests that MGE abundance is less likely to change due to direct accumulation by introduced invaders. Therefore, the change in prophage abundance under E-A treatment is expected due to the more extensive input derived from invasive communities with a significantly higher quantity of prophages than other invasive communities and the resident community (Supplementary Figures S1 and S9). The introduced prophages were viable and capable of lysing their hosts and reproducing within the coalescent community during environmental changes [39]. In this case, the coalescence process could be the source of various stresses for invaders, such as nutritional competitions between invaders and residents, antibiotics produced by residents, and quorum sensing derived from residents, thereby leading to phage excision.

Alternatively, the changes in the total abundance of other MGEs, such as plasmids, ICE, and IS, may primarily rely on shifts in microbial communities, including establishments of specific invaders and the thriving or suppression of particular resident taxa. Importantly, we observed a common effect across treatments: the significant increase in the total abundance of IS after coalescence. This phenomenon is supported by the increased abundance of “IE division” genes associated with integration and excision, which coincides with the function of the ISs (Brown et al., 2022). Additionally, the significant negative correlation between the abundance of ISs in invasive and coalescent communities on Day 5 suggests that ISs did not directly contribute to increased ISs in coalescent communities (Supplementary Figures S9 and S10). Moreover, the abundance of ISs enriched under each treatment related to uninvaded control is significantly lower in invasive communities than in uninvaded control and coalescent communities, let alone the low invasion rate (approximately 5% of the density of resident taxa) (Supplementary Figure S14) we applied. Our results suggest that the increase in IS abundance is an intrinsic response of the resident community following coalescence.

### Competitive stress triggers changes in the MGE profile after coalescence

Competition for resources between invaders and residents is a common interaction after coalescence. Our results suggest that the response of MGEs is significantly correlated to the probability of such competition between invaders and residents. Two noteworthy phenomena emerged from this study: First, positive correlations were predominantly observed on Day 5, suggesting higher competitive stress at the early coalescence stage. This aligns with our previous observations that the survival rate of invaders was lower, and the impact on community composition was less severe at the late stages of the coalescence (Liu and Salles, 2023). Second, relatively stronger correlations were found with carbon-cycling genes compared to nitrogen, phosphorus, and sulfur-related genes, indicating a more intensive competition for carbon sources.

These two points are further supported by network analyses of potential association patterns between MGEs and genes related to microbial antibiotic resistance and nutrient cycling. MGE-mediated transfers of antibiotic resistance genes and resource utilization genes are the main ways microorganisms enhance their fitness (Toft and Andersson, 2010). Especially in terms of harnessing energy, microorganisms may have the opportunity to utilize novel carbon sources after acquiring genes from other microorganisms through HGT, thereby gaining survival advantages in resource-limited or contaminated environments (Jain et al., 2003). The network analyses suggest that MGEs exhibited more intensive and positive correlations with these genes on Day 5 than Day 60, indicating the more intensive adaptive pattern of coalescent communities at the early stage after coalescence. Furthermore, within these associations, genes involved in nutrient cycling, especially carbon-cycling genes, appeared more susceptible to MGE influences on Day 5. These findings underscore the significance of invader-triggered competitions and nutrient-cycling genes for communities responding to such disturbances.

### The critical role of ISs in HGTs

Our findings support the notion that the increased abundance of ISs is likely a direct outcome of horizontal gene transfers (HGTs) within the community by evaluating the neighbors of intermediate contigs containing IS fragments. Given that IS fragments are relatively short in length (typically 0.7–2.5 kb long (Siguier et al., 2014)), they are more likely to be assembled into a single contig in the assembly graph, enabling a more accurate analysis of the number of contig neighbors to represent the frequency of ISs in different genomes at the community level. The dissemination of IS fragments occurs through multiple mechanisms, including vectors like plasmids via conjugation or through their host genomes via replication and translocation (Preston et al., 2004). This can lead to repetitive sequences in the meta-genome graph. Our results show that the increased abundance of ISs was significantly positively correlated with the increased contig connectivity in the assembly graph, and the coefficient was higher for ISs than other MGEs such as ICEs, plasmids, and integrons.

The Network analysis unveiled notable associations between ISs and genes related to nutrient cycling and antibiotic resistance. Interestingly, the proportion of IS-related links in the network increased on Day 60, suggesting a potentially more pronounced role of ISs at this stage. This increase in IS-related links was accompanied by a decrease in the proportion of links associated with other MGEs. The enhanced connectivity of ISs with nutrient-cycling and antibiotic genes underscores their potential role in shaping community functions and adaptation. The association of ISs with nutrient-cycling genes highlights their relevance in facilitating resource utilization and metabolic versatility within the coalescent communities. Furthermore, the links between ISs and antibiotic genes indicate their potential involvement in disseminating antibiotic resistance traits, which can enhance the survival and competitiveness of microorganisms in resource-limited or contaminated environments. However, the insertion sites of ISs may be approximately random, and the transfer may activate or inactivate nearby functional genes (Vandecraen et al., 2017). In addition, the fragments transferred by ISs do not necessarily result in better host fitness (Vandecraen et al., 2017). Therefore, the genetic diversification suggested by increased IS abundance does not necessarily imply better fitness or adaptivity of microbial communities. Indeed, we found that, on Day 60, a large proportion of ISs that enriched on Day 5 was not varied from the uninvaded control, and the total abundance of ISs was recovered to a similar level across treatments. These reflect that ISs can promote and constrain microbial mutation and adaptivity (Consuegra et al., 2021). Nevertheless, on Day 60, the network analysis ISs displayed more connections with potential hosts, represented by ASVs. These observations suggest that a portion of ISs carrying adaptive genes via horizontal gene transfer may facilitate the acquisition of adaptations by the host and establish a broader presence within different hosts at the late stage of coalescence. Overall, our results highlight the importance of ISs in community genetic diversity and adaptation following coalescence.

It is worth noting that further investigations are needed to unravel the specific mechanisms underlying the interactions between ISs and host genomes, as well as the functional implications of these associations. To achieve this, long-length sequencing techniques can provide a more accurate assessment of the relationships between IS fragments and adjacent functional elements and their respective host genomes (Zhou et al., 2020; Che et al., 2021). Besides, by adopting targeted metagenomic sequencing approaches (Dunon et al., 2018), it becomes easier to track the dynamic patterns of specific ISs throughout the process of community succession in the context of community coalescence. This targeted approach is a powerful tool for characterizing the precise role of ISs in driving community dynamics and adaptation.

## Conclusions

This study provided insights into the evolutionary perspective of soil microbial communities under the coalescence scenario. On the one hand, our results confirm that competition for resources could be the primary mechanism underlying coalescences in soil. This has implications for understanding soil microbial community dynamics and regulating inoculating beneficial consortia in agroecosystems. Specifically, predicting the consequences of community coalescences requires consideration of a lose-lose competitive situation (Liu and Salles, 2023), and in agricultural management, taking measures to reduce competition for inoculum will increase inoculation benefits (Liu et al., 2022). On the other hand, our results suggest a rapid response of MGE profiles and increased HGT potential triggered by coalescence, where ISs may play a vital role in promoting genomic diversification in soil communities. The artificially manipulated and improved adaptation of microbial communities to drought, heat, or cold stress may enhance our ability to predict and manage the adaptability of ecosystems to changing climates (Allsup et al., 2023). Although our results are limited to profile functional traits at the community level, they point to a need for further investigation of specific evolution within coalescent communities using new technologies such as third-generation sequencing and the coalescence impact on community fitness against future disturbances and subsequent ecosystem functions.

## Acknowledgments

This work was financed by the ERA-NET Cofund SusCrop project potatoMETAbiome (Grant No 771134). Xipeng Liu was supported by a scholarship from the China Scholarship Council. The authors acknowledge the Center for Information Technology at the University of Groningen for providing access to the Peregrine high-performance computing cluster.

## Availability of data and materials

All the raw sequencing data were deposited in the National Center for Biotechnology Information Sequence Read Archive under the accession number PRJNA843110.

## References

1. Alcock, B.P., Raphenya, A.R., Lau, T.T.Y., … G.V., McArthur, A.G., 2020. CARD 2020: antibiotic resistome surveillance with the comprehensive antibiotic resistance database. Nucleic Acids Research 48, D517–D525. doi:10.1093/nar/gkz935

2. Allsup, C.M., George, I., Lankau, R.A., 2023. Shifting microbial communities can enhance tree tolerance to changing climates. Science 380, 835–840. doi:10.1126/science.adf2027

3. Arndt, D., Grant, J.R., Marcu, A., Sajed, T., Pon, A., Liang, Y., Wishart, D.S., 2016. PHASTER: a better, faster version of the PHAST phage search tool. Nucleic Acids Research 44, W16–21. doi:10.1093/nar/gkw387

4. Arnold, B.J., Huang, I.-T., Hanage, W.P., 2022. Horizontal gene transfer and adaptive evolution in bacteria. Nature Reviews Microbiology 20, 206–218. doi:10.1038/s41579-021-00650-4

5. Bastian, M., Heymann, S., Jacomy, M., 2009. Gephi: An Open Source Software for Exploring and Manipulating Networks. Proceedings of the International AAAI Conference on Web and Social Media 3, 361–362. doi:10.1609/icwsm.v3i1.13937

6. Brito, I.L., 2021. Examining horizontal gene transfer in microbial communities. Nature Reviews Microbiology 19, 442–453. doi:10.1038/s41579-021-00534-7

7. Brown, C.L., Mullet, J., Hindi, F., Stoll, J.E., Gupta, S., Choi, M., Keenum, I., Vikesland, P., Pruden, A., Zhang, L., 2022. mobileOG-db: a Manually Curated Database of Protein Families Mediating the Life Cycle of Bacterial Mobile Genetic Elements. Applied and Environmental Microbiology 88, e00991–22. doi:10.1128/aem.00991-22

8. Buchfink, B., Reuter, K., Drost, H.-G., 2021. Sensitive protein alignments at tree-of-life scale using DIAMOND. Nature Methods 18, 366–368. doi:10.1038/s41592-021-01101-x

9. Cantarel, B.L., Coutinho, P.M., Rancurel, C., Bernard, T., Lombard, V., Henrissat, B., 2009. The Carbohydrate-Active EnZymes database (CAZy): an expert resource for Glycogenomics. Nucleic Acids Research 37, D233–D238. doi:10.1093/nar/gkn663

10. Chase, A.B., Weihe, C., Martiny, J.B.H., 2021. Adaptive differentiation and rapid evolution of a soil bacterium along a climate gradient. Proceedings of the National Academy of Sciences 118, e2101254118. doi:10.1073/pnas.2101254118

11. Che, Y., Yang, Y., Xu, X., Břinda, K., Polz, M.F., Hanage, W.P., Zhang, T., 2021. Conjugative plasmids interact with insertion sequences to shape the horizontal transfer of antimicrobial resistance genes. Proceedings of the National Academy of Sciences 118, e2008731118. doi:10.1073/pnas.2008731118

12. Chen, Yuxin, Chen, Yongsheng, Shi, C., Huang, Z., Zhang, Y., Li, S., Li, Y., Ye, J., Yu, C., Li, Z., Zhang, X., Wang, J., Yang, H., Fang, L., Chen, Q., 2018. SOAPnuke: a MapReduce acceleration-supported software for integrated quality control and preprocessing of high-throughput sequencing data. GigaScience 7, 1–6. doi:10.1093/gigascience/gix120

13. Consuegra, J., Gaffé, J., Lenski, R.E., Hindré, T., Barrick, J.E., Tenaillon, O., Schneider, D., 2021. Insertion-sequence-mediated mutations both promote and constrain evolvability during a long-term experiment with bacteria. Nature Communications 12, 980. doi:10.1038/s41467-021-21210-7

14. Diaz-Colunga, J., Lu, N., Sanchez-Gorostiaga, A., Chang, C.-Y., Cai, H.S., Goldford, J.E., Tikhonov, M., Sánchez, Á., 2022. Top-down and bottom-up cohesiveness in microbial community coalescence. Proceedings of the National Academy of Sciences 119. doi:10.1073/pnas.2111261119

15. Douglas, G.M., Langille, M.G.I., 2019. Current and Promising Approaches to Identify Horizontal Gene Transfer Events in Metagenomes. Genome Biology and Evolution 11, 2750–2766. doi:10.1093/gbe/evz184

16. Dunon, V., Bers, K., Lavigne, R., Top, E.M., Springael, D., 2018. Targeted metagenomics demonstrates the ecological role of IS1071 in bacterial community adaptation to pesticide degradation. Environmental Microbiology 20, 4091–4111. doi:10.1111/1462-2920.14404

17. Galata, V., Fehlmann, T., Backes, C., Keller, A., 2019. PLSDB: a resource of complete bacterial plasmids. Nucleic Acids Research 47, D195–D202. doi:10.1093/nar/gky1050

18. Hawkins, N.J., Bass, C., Dixon, A., Neve, P., 2019. The evolutionary origins of pesticide resistance. Biological Reviews 94, 135–155. doi:10.1111/brv.12440

19. Horne, T., Orr, V.T., Hall, J.P., 2023. How do interactions between mobile genetic elements affect horizontal gene transfer? Current Opinion in Microbiology 73, 102282. doi:10.1016/j.mib.2023.102282

20. Huet, S., Romdhane, S., Breuil, M.-C., Bru, D., Mounier, A., Spor, A., Philippot, L., 2023. Experimental community coalescence sheds light on microbial interactions in soil and restores impaired functions. Microbiome 11, 42. doi:10.1186/s40168-023-01480-7

21. Hyatt, D., Chen, G.-L., LoCascio, P.F., Land, M.L., Larimer, F.W., Hauser, L.J., 2010. Prodigal: prokaryotic gene recognition and translation initiation site identification. BMC Bioinformatics 11, 119. doi:10.1186/1471-2105-11-119

22. Jain, R., Rivera, M.C., Moore, J.E., Lake, J.A., 2003. Horizontal Gene Transfer Accelerates Genome Innovation and Evolution. Molecular Biology and Evolution 20, 1598–1602. doi:10.1093/molbev/msg154

23. Li, D., Liu, C.-M., Luo, R., Sadakane, K., Lam, T.-W., 2015. MEGAHIT: an ultra-fast single-node solution for large and complex metagenomics assembly via succinct de Bruijn graph. Bioinformatics 31, 1674–1676. doi:10.1093/bioinformatics/btv033

24. Liu, M., Li, X., Xie, Y., Bi, D., Sun, J., Li, J., Tai, C., Deng, Z., Ou, H.-Y., 2019. ICEberg 2.0: an updated database of bacterial integrative and conjugative elements. Nucleic Acids Research 47, D660–D665. doi:10.1093/nar/gky1123

25. Liu, X., Mei, S., Salles, J.F., 2023. Do inoculated microbial consortia perform better than single strains in living soil? A meta-analysis. doi:10.1101/2023.03.17.533112

26. Liu, X., Roux, X.L., Salles, J.F., 2022. The Legacy of Microbial Inoculants in Agroecosystems and Potential for Tackling Climate Change Challenges. IScience 0. doi:10.1016/j.isci.2022.103821

27. Liu, X., Salles, J., 2023. Lose-lose consequences of bacterial community-driven invasions in soil. doi:10.21203/rs.3.rs-2506521/v1

28. Moura, A., Soares, M., Pereira, C., Leitão, N., Henriques, I., Correia, A., 2009. INTEGRALL: a database and search engine for integrons, integrases and gene cassettes. Bioinformatics 25, 1096–1098. doi:10.1093/bioinformatics/btp105

29. Pärnänen, K.M.M., Narciso-da-Rocha, C., Kneis, D., Berendonk, T.U., Cacace, D., Do, T.T., Elpers, C., Fatta-Kassinos, D., Henriques, I., Jaeger, T., Karkman, A., Martinez, J.L., Michael, S.G., Michael-Kordatou, I., O’Sullivan, K., Rodriguez-Mozaz, S., Schwartz, T., Sheng, H., Sørum, H., Stedtfeld, R.D., Tiedje, J.M., Giustina, S.V.D., Walsh, F., Vaz-Moreira, I., Virta, M., Manaia, C.M., 2019. Antibiotic resistance in European wastewater treatment plants mirrors the pattern of clinical antibiotic resistance prevalence. Science Advances 5, eaau9124. doi:10.1126/sciadv.aau9124

30. Pasternak, C., Ton-Hoang, B., Coste, G., Bailone, A., Chandler, M., Sommer, S., 2010. Irradiation-Induced Deinococcus radiodurans Genome Fragmentation Triggers Transposition of a Single Resident Insertion Sequence. PLOS Genetics 6, e1000799. doi:10.1371/journal.pgen.1000799

31. Patro, R., Duggal, G., Love, M.I., Irizarry, R.A., Kingsford, C., 2017. Salmon provides fast and bias-aware quantification of transcript expression. Nature Methods 14, 417–419. doi:10.1038/nmeth.4197

32. Preston, A., Parkhill, J., Maskell, D.J., 2004. The Bordetellae: lessons from genomics. Nature Reviews Microbiology 2, 379–390. doi:10.1038/nrmicro886

33. Ramoneda, J., Le Roux, J., Stadelmann, S., Frossard, E., Frey, B., Gamper, H.A., 2021. Soil microbial community coalescence and fertilization interact to drive the functioning of the legume–rhizobium symbiosis. Journal of Applied Ecology 58, 2590–2602. doi:10.1111/1365-2664.13995

34. Rillig, M.C., Antonovics, J., Caruso, T., Lehmann, A., Powell, J.R., Veresoglou, S.D., Verbruggen, E., 2015. Interchange of entire communities: microbial community coalescence. Trends in Ecology & Evolution 30, 470–476. doi:10.1016/j.tree.2015.06.004

35. Rillig, M.C., Lehmann, A., Aguilar-Trigueros, C.A., Antonovics, J., Caruso, T., Hempel, S., Lehmann, J., Valyi, K., Verbruggen, E., Veresoglou, S.D., Powell, J.R., 2016. Soil microbes and community coalescence. Pedobiologia 59, 37–40. doi:10.1016/j.pedobi.2016.01.001

36. Siguier, P., Gourbeyre, E., Chandler, M., 2014. Bacterial insertion sequences: their genomic impact and diversity. FEMS Microbiology Reviews 38, 865–891. doi:10.1111/1574-6976.12067

37. Siguier, P., Perochon, J., Lestrade, L., Mahillon, J., Chandler, M., 2006. ISfinder: the reference centre for bacterial insertion sequences. Nucleic Acids Research 34, D32–36. doi:10.1093/nar/gkj014

38. Steinegger, M., Söding, J., 2017. MMseqs2 enables sensitive protein sequence searching for the analysis of massive data sets. Nature Biotechnology 35, 1026–1028. doi:10.1038/nbt.3988

39. Toft, C., Andersson, S.G.E., 2010. Evolutionary microbial genomics: insights into bacterial host adaptation. Nature Reviews Genetics 11, 465–475. doi:10.1038/nrg2798

40. Tu, Q., Lin, L., Cheng, L., Deng, Y., He, Z., 2019. NCycDB: a curated integrative database for fast and accurate metagenomic profiling of nitrogen cycling genes. Bioinformatics 35, 1040–1048. doi:10.1093/bioinformatics/bty741

41. Vandecraen, J., Chandler, M., Aertsen, A., Van Houdt, R., 2017. The impact of insertion sequences on bacterial genome plasticity and adaptability. Critical Reviews in Microbiology 43, 709–730. doi:10.1080/1040841X.2017.1303661

42. Wang, M., Goh, Y.-X., Tai, C., Wang, H., Deng, Z., Ou, H.-Y., 2022. VRprofile2: detection of antibiotic resistance-associated mobilome in bacterial pathogens. Nucleic Acids Research 50, W768–W773. doi:10.1093/nar/gkac321

43. Woods, L.C., Gorrell, R.J., Taylor, F., Connallon, T., Kwok, T., McDonald, M.J., 2020. Horizontal gene transfer potentiates adaptation by reducing selective constraints on the spread of genetic variation. Proceedings of the National Academy of Sciences 117, 26868–26875. doi:10.1073/pnas.2005331117

44. Yao, Y., Maddamsetti, R., Weiss, A., Ha, Y., Wang, T., Wang, S., You, L., 2022. Intra– and interpopulation transposition of mobile genetic elements driven by antibiotic selection. Nature Ecology & Evolution 6, 555–564. doi:10.1038/s41559-022-01705-2

45. Yu, X., Zhou, J., Song, W., Xu, M., He, Q., Peng, Y., Tian, Y., Wang, C., Shu, L., Wang, S., Yan, Q., Liu, J., Tu, Q., He, Z., 2021. SCycDB: A curated functional gene database for metagenomic profiling of sulphur cycling pathways. Molecular Ecology Resources 21, 924–940. doi:10.1111/1755-0998.13306

46. Zeng, J., Tu, Q., Yu, X., Qian, L., Wang, C., Shu, L., Liu, F., Liu, S., Huang, Z., He, J., Yan, Q., He, Z., 2022. PCycDB: a comprehensive and accurate database for fast analysis of phosphorus cycling genes. Microbiome 10, 101. doi:10.1186/s40168-022-01292-1

47. Zhou, W., Emery, S.B., Flasch, D.A., Wang, Y., Kwan, K.Y., Kidd, J.M., Moran, J.V., Mills, R.E., 2020. Identification and characterization of occult human-specific LINE-1 insertions using long-read sequencing technology. Nucleic Acids Research 48, 1146– 1163. doi:10.1093/nar/gkz1173

