## Supplemental Materials for "Community coalescence altered the potential of horizontal gene transfers within the native soil microbiome"

**Supplementary information**

The Supplementary information includes Supplementary Figure S1-14 and Table S1.

***Supplementary figures***

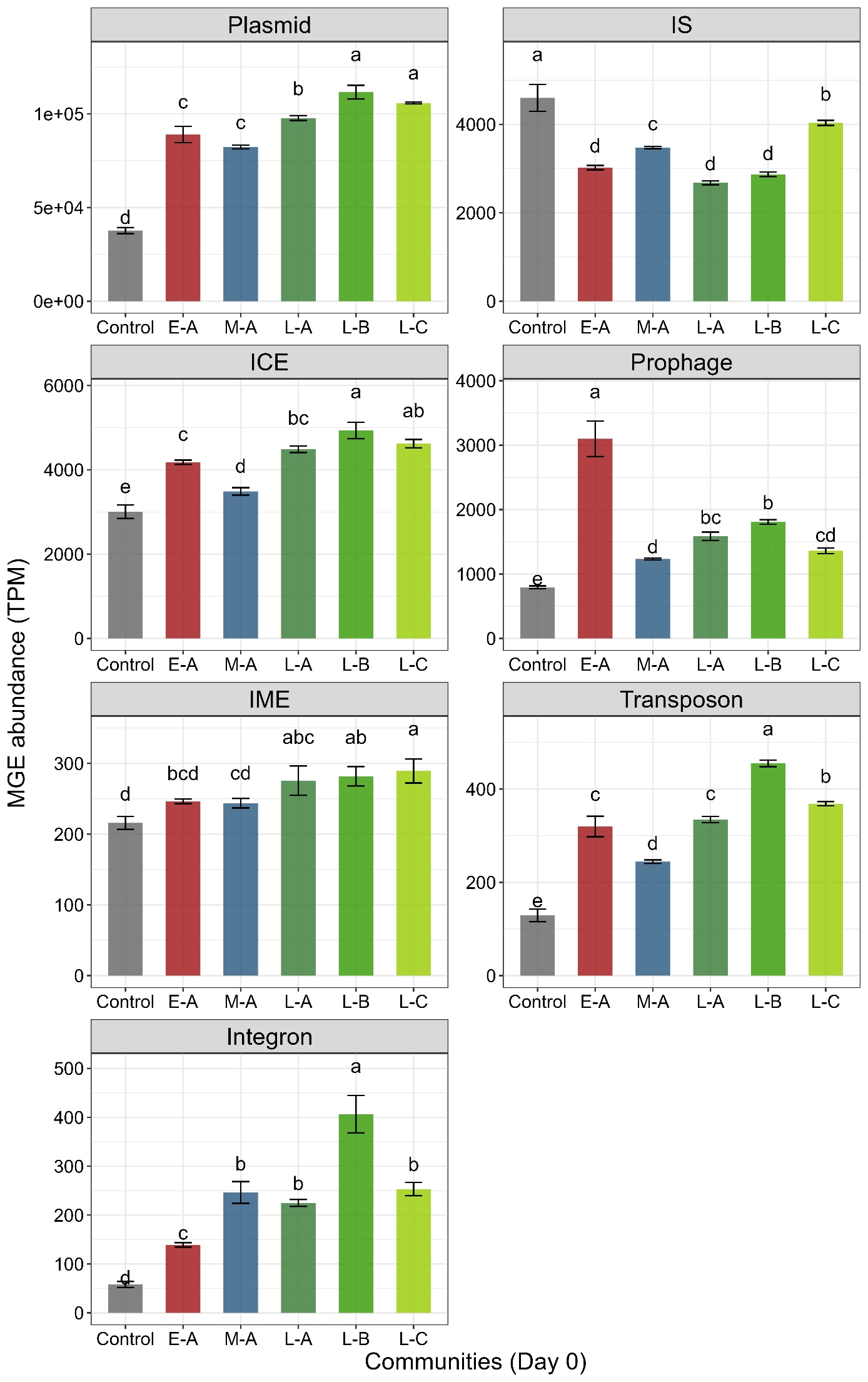

Figure S1. The abundance of MGEs in original (Control) and invasive communities. The letters above bars indicate the difference among treatments with the Tukey test (*p* < 0.05).

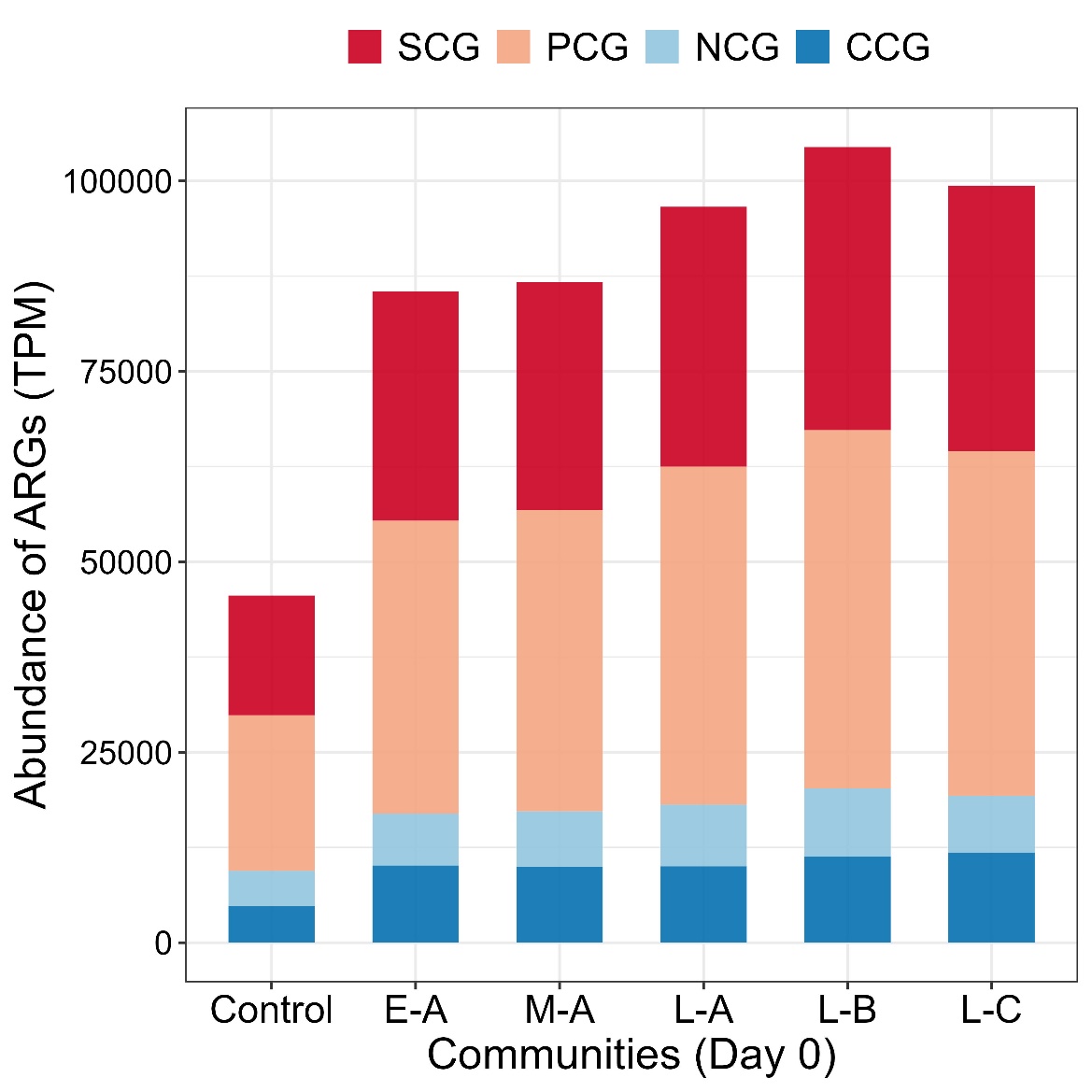

Figure S2. The abundance of CNPS-cycling genes in original (Control) and invasive communities on Day 0. CCGs, carbon-cycling genes; NCGs, nitrogen-cycling genes; PCGs, phosphorus-cycling genes; and SCGs, sulfur-cycling genes.

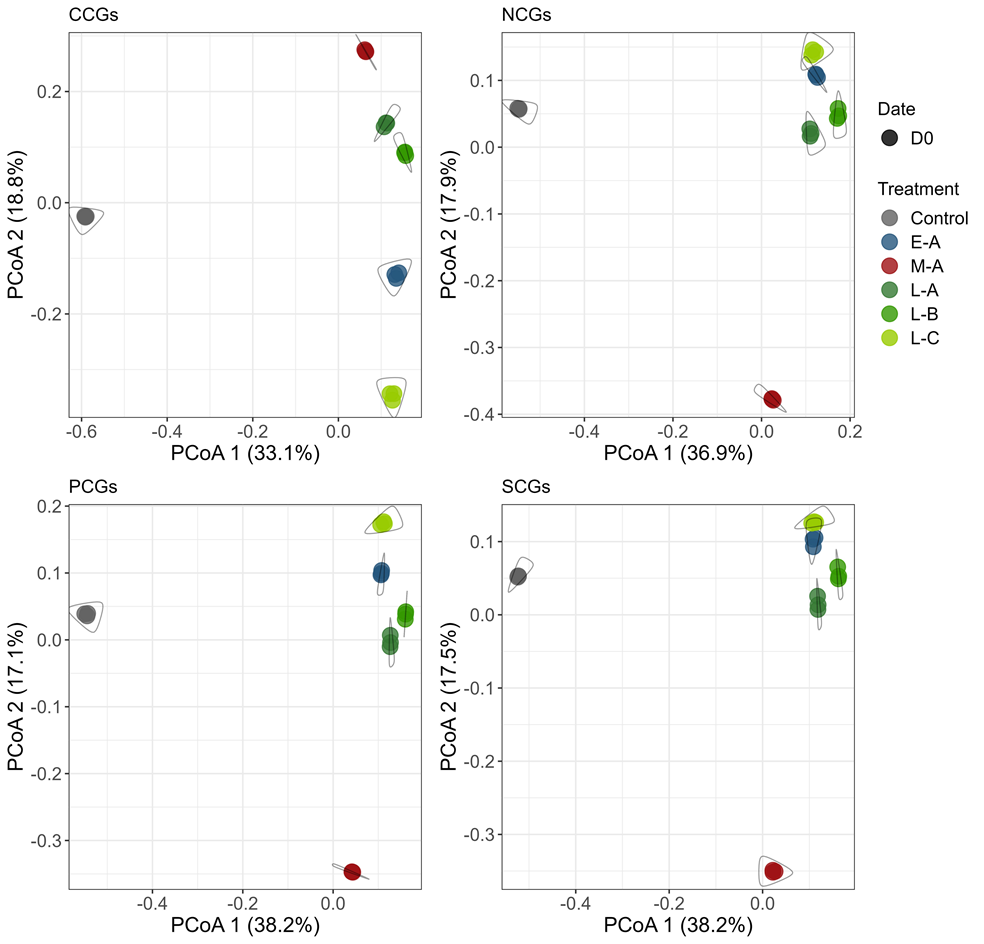

Figure S3. Principal Coordinates Analysis (PCoA) on profiles of CNPS-cycling genes in original (Control) and invasive communities on Day 0. CCGs, carbon-cycling genes; NCGs, nitrogen-cycling genes; PCGs, phosphorus-cycling genes; and SCGs, sulfur-cycling genes.

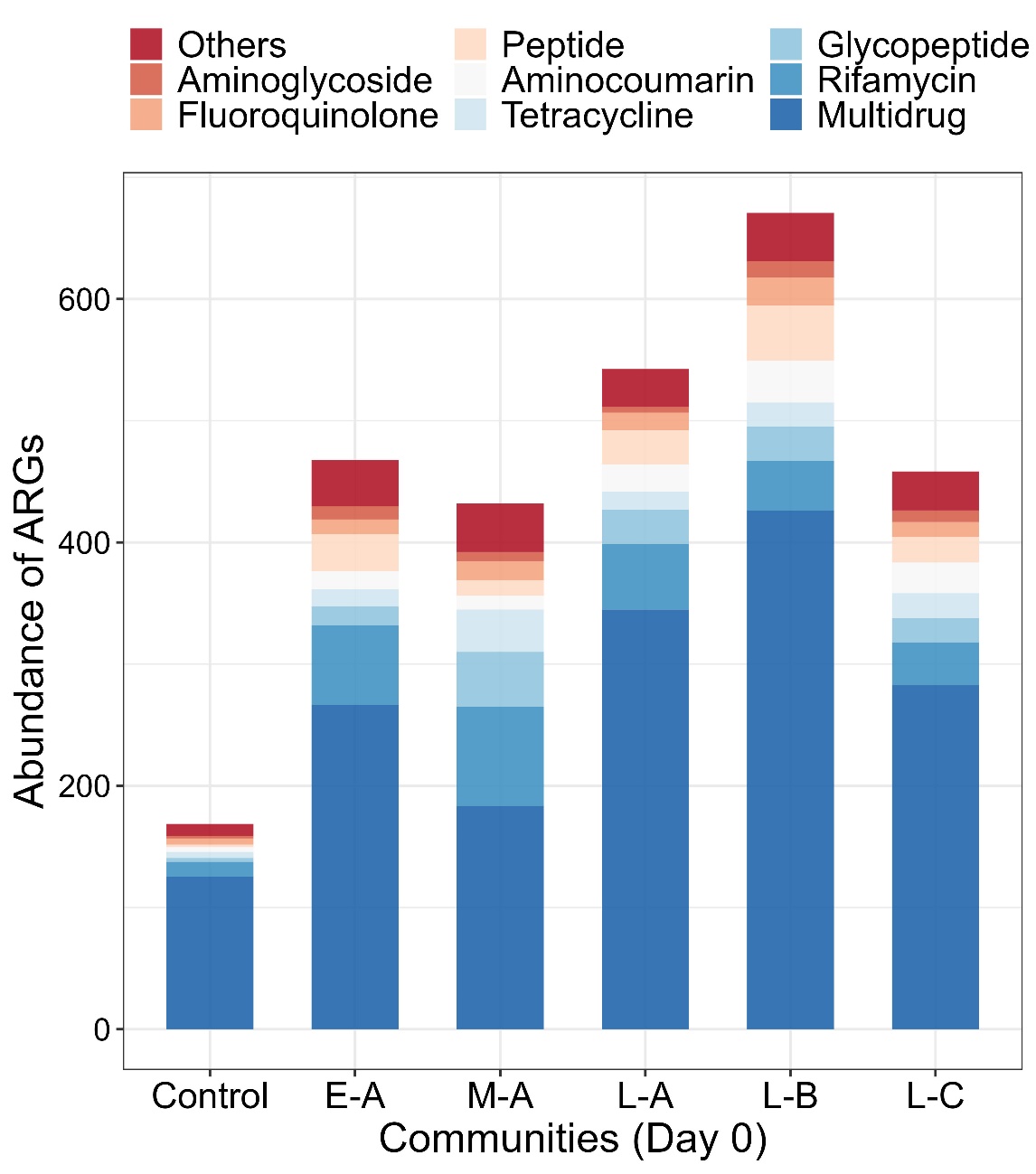

Figure S4. The abundance of ARGs in original (Control) and invasive communities on Day 0. The ARGs were classified by drug types.

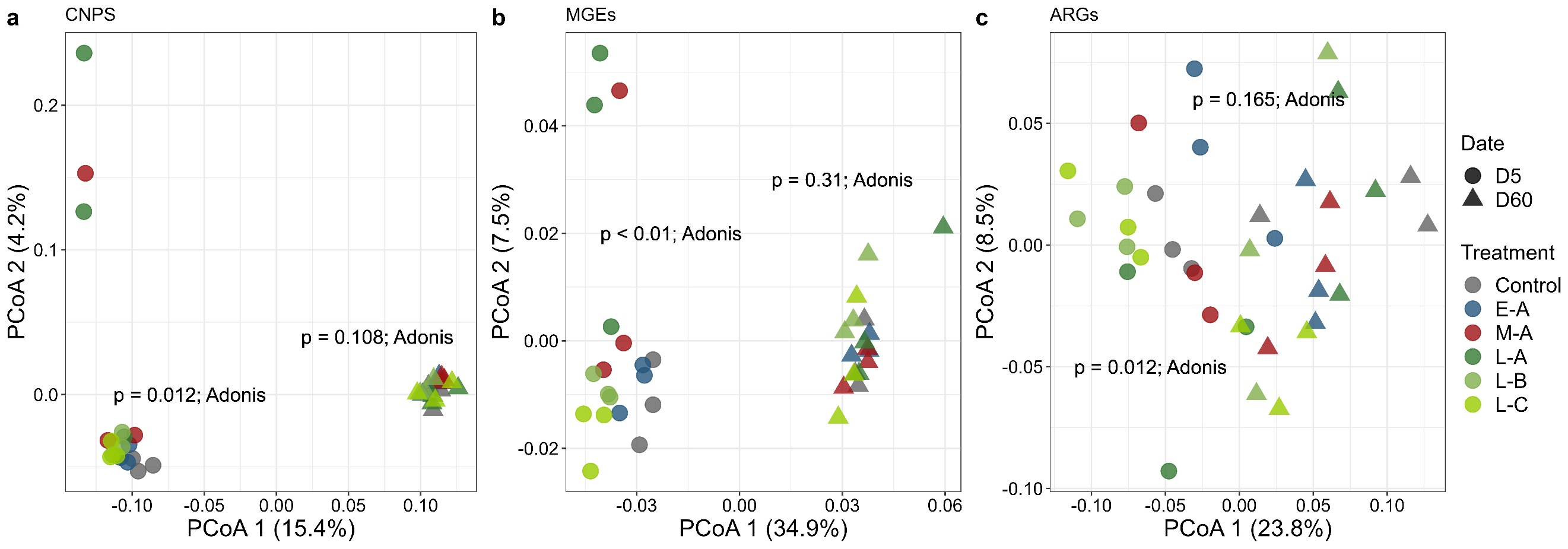

Figure S5. Principal Coordinates Analysis (PCoA) on the effect of community coalescence on the functional traits (CNPS-cycling genes, MGEs, and ARGs). The p-value of Permutational Multivariate Analysis of Variance (Adonis) close to different points indicated the functional variance for each date.

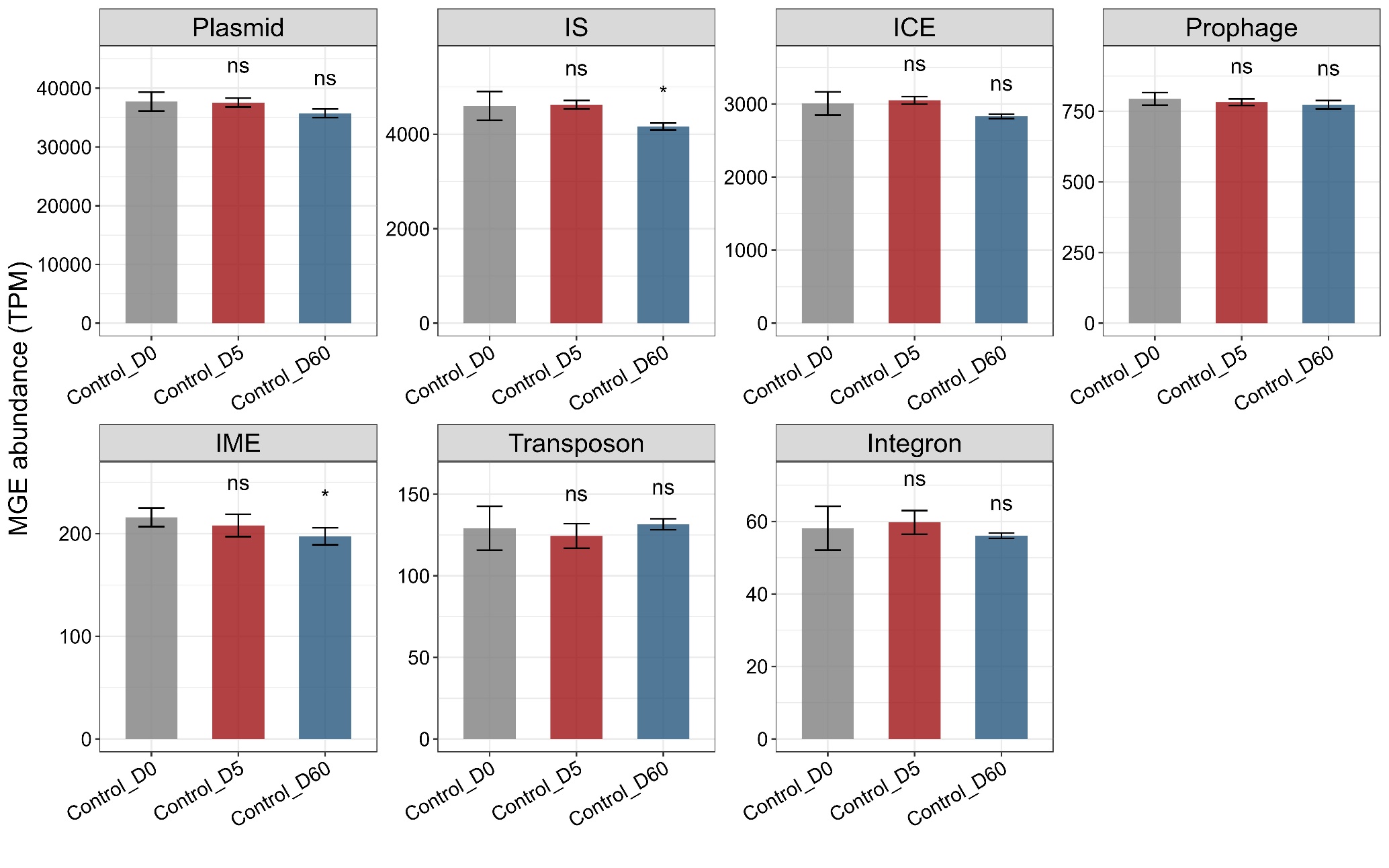
Figure S6. MGE abundance in the uninvaded soils across time. D0, D5, and D60 represent before the invasion, five days, and 60 days after the invasion. Asterisks suggest the significant difference in MGEs’ abundance between control_D0 and other samples with a t-test (*, *p* < 0.05).

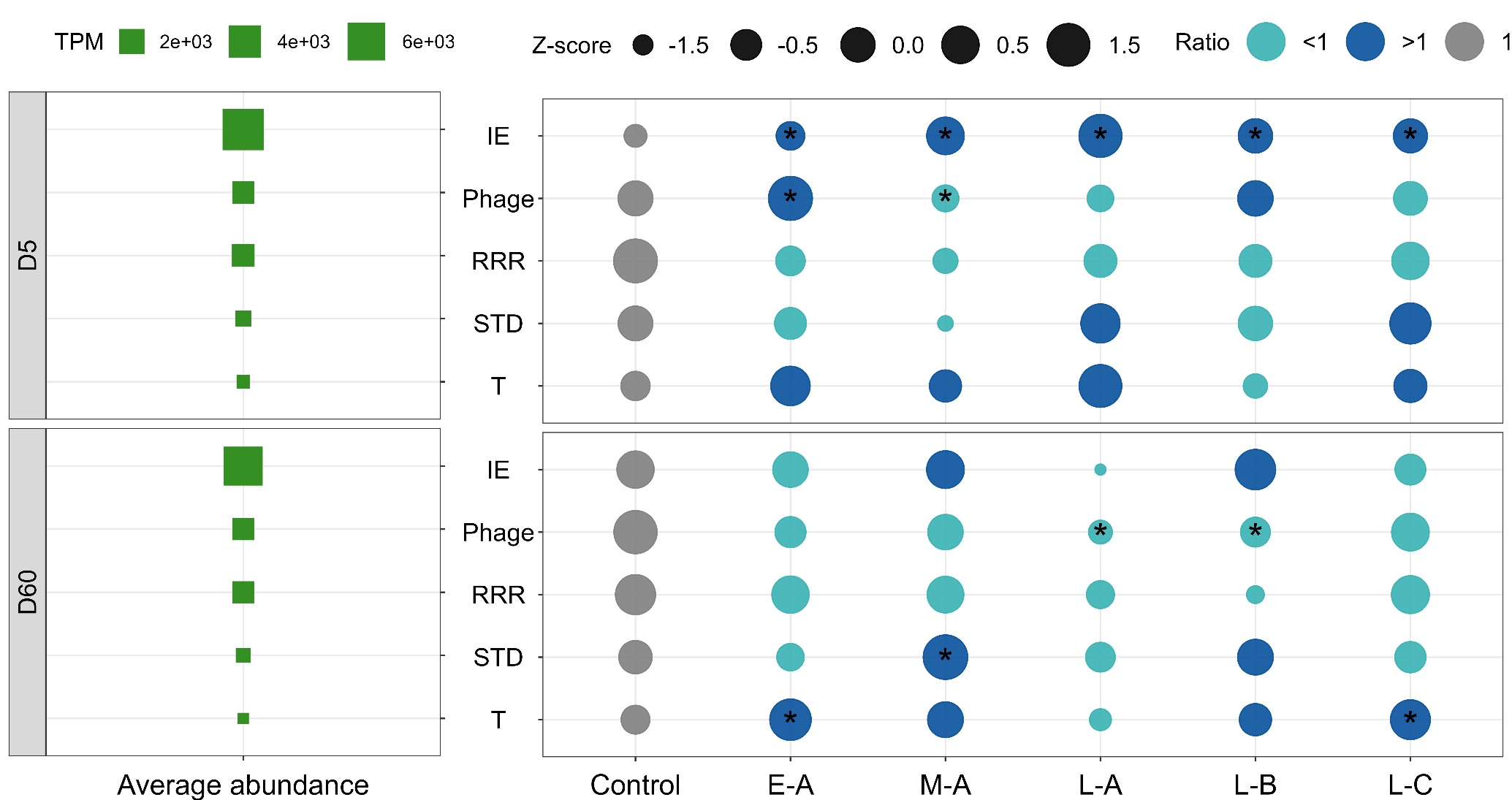

Figure S7. The functional classification of MGE genes annotated against the MobileOG database. The five functional divisions are integration and excision (IE), phage-related process (P), replication/recombination/repair (RRR), stable/transfer/defense (STD), and Transfer (T). The abundance (TPM) of MGEs was transformed with Z-score. The color indicates the ratio of MGE abundance in relation to the uninvaded control. Asterisks suggest the significant difference in MGEs’ abundance between uninvaded control and coalescent communities with a t-test (*p* < 0.05).

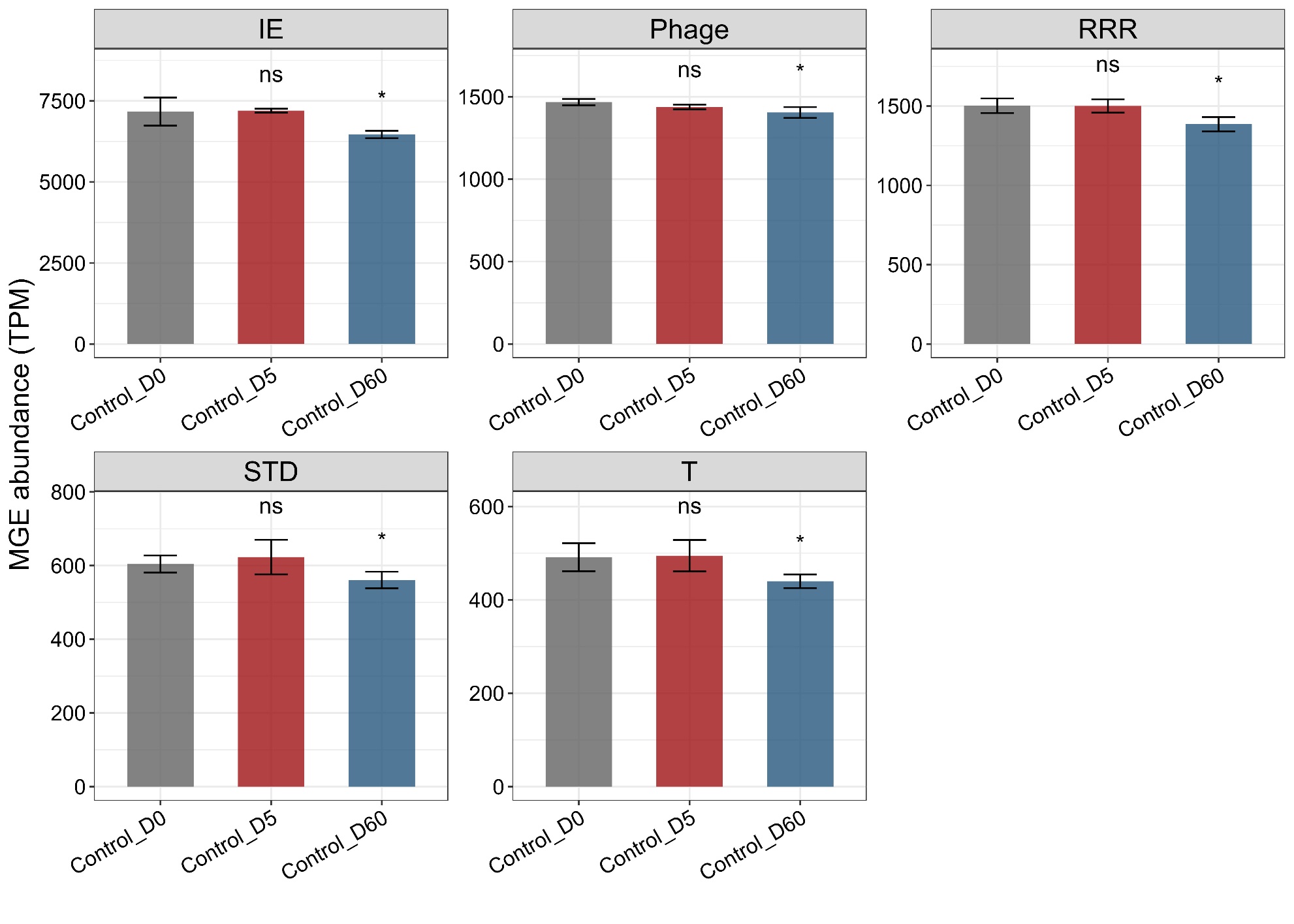
Figure S8. MGE abundance (annotated against the MobileOG database) in the uninvaded soils across time. D0, D5, and D60 represent before invasion, five days, and 60 days after the invasion. Asterisks suggest the significant difference in MGEs’ abundance between control_D0 and other samples with a t-test (*, *p* < 0.05).

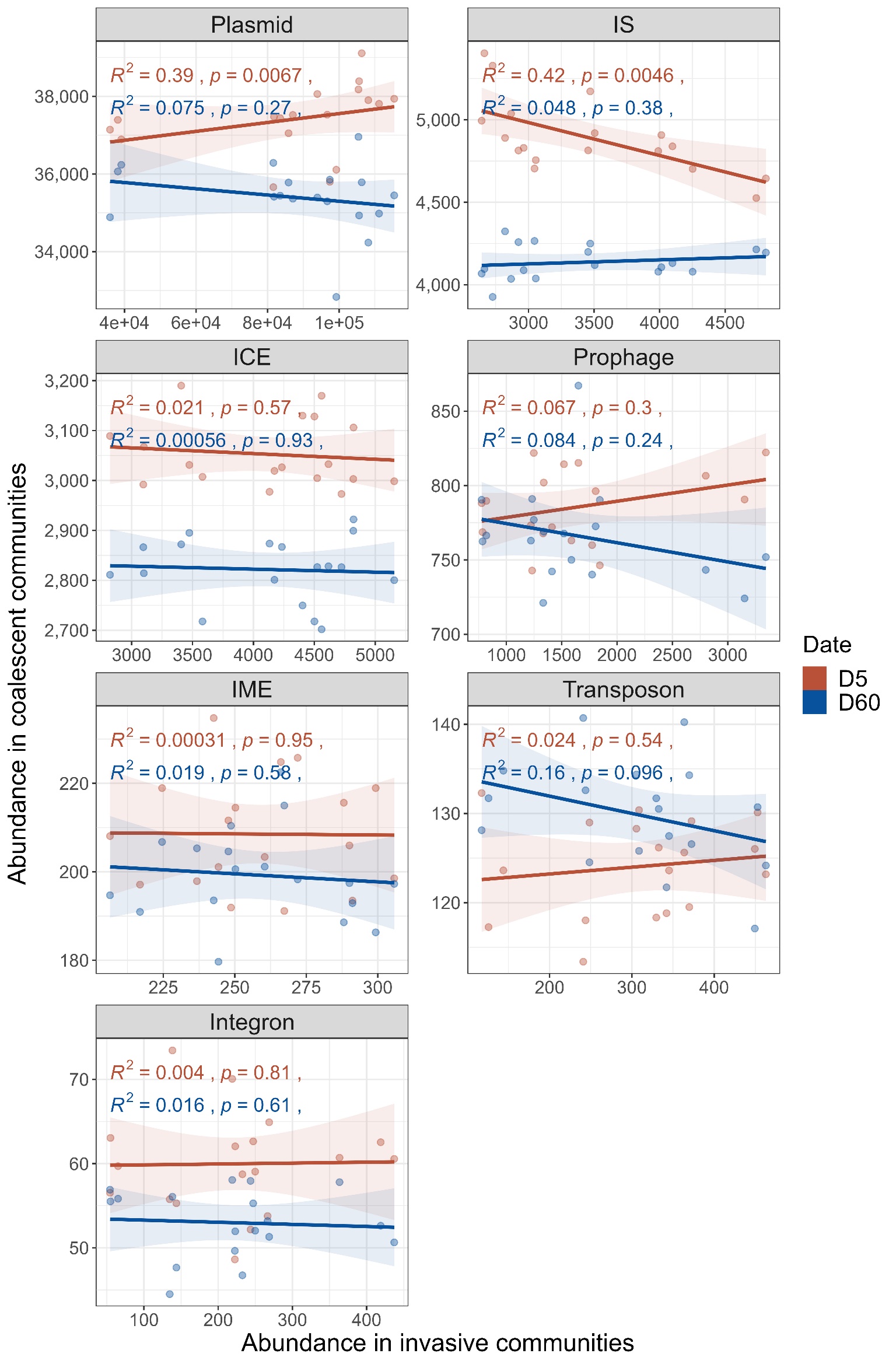

Figure S9. Spearman’s correlation between the abundance of MGE gene (classified by types) in invasive and coalescent communities.

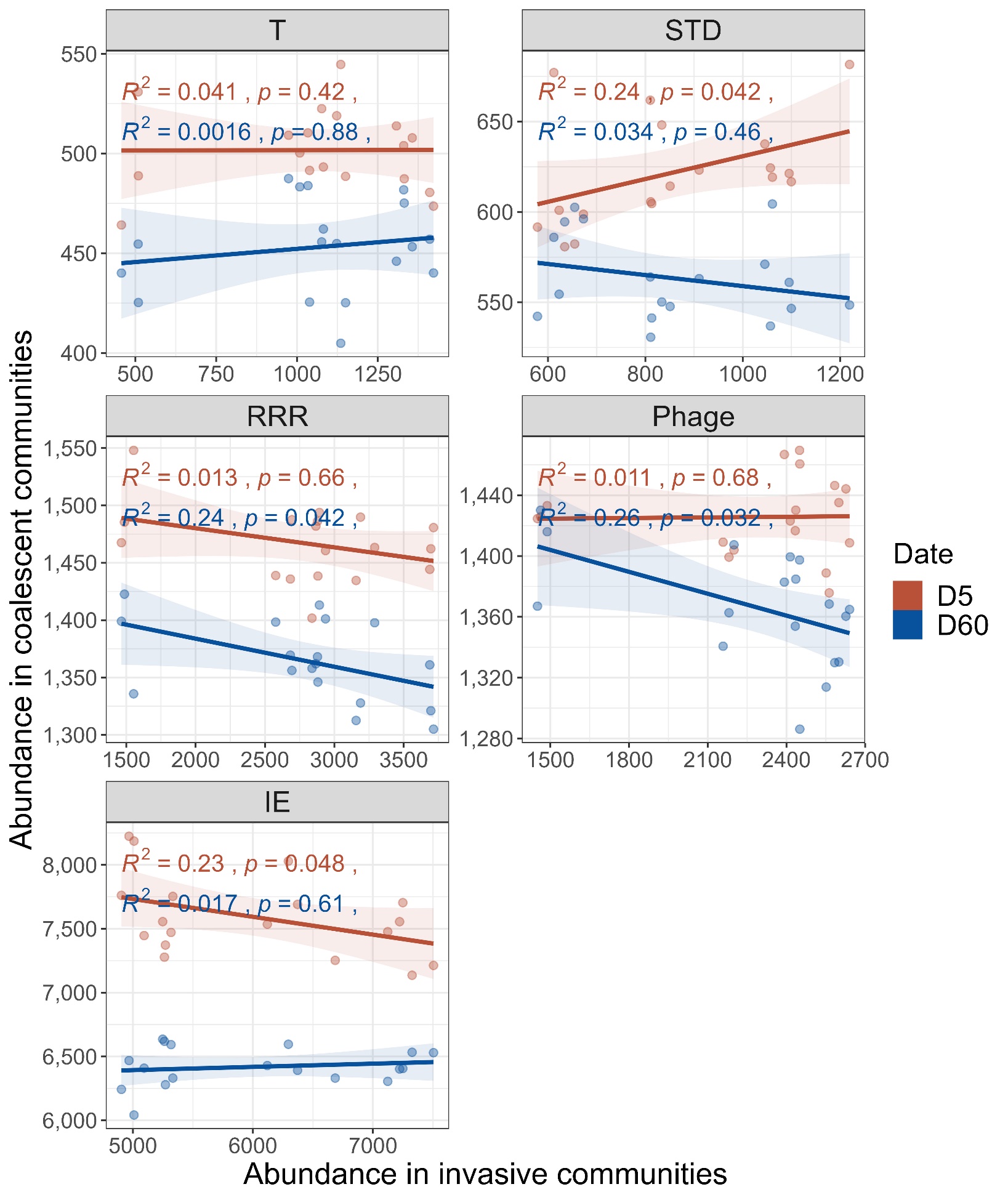

Figure S10. Spearman’s correlation between the abundance of MGE gene (classified by functions) in invasive and coalescent communities. The MGE genes are categorized into five functional divisions: integration and excision (IE), phage-related processes (Phage), replication/recombination/repair (RRR), stable/transfer/defense (STD), and transfer (T).

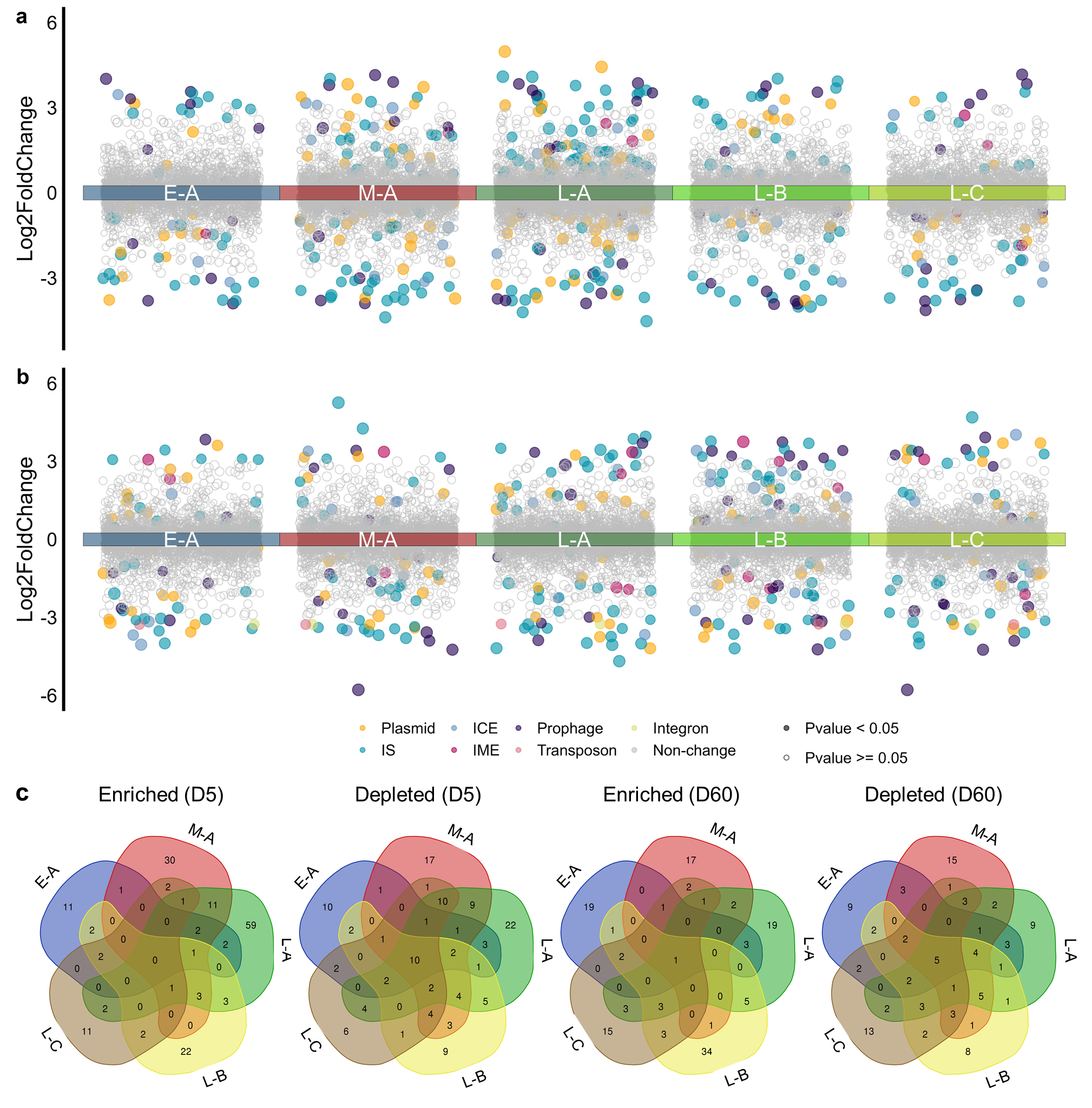

Figure S11. The abundance changes of MGEs at the gene family level compared to the uninvaded control. The variance was estimated using edgeR, and the significance was *p* < 0.05. (a) and (b) represent the data for Day 5 and Day 60, respectively. (c) The Venn diagram shows MGE types with significantly changed abundance across treatments.

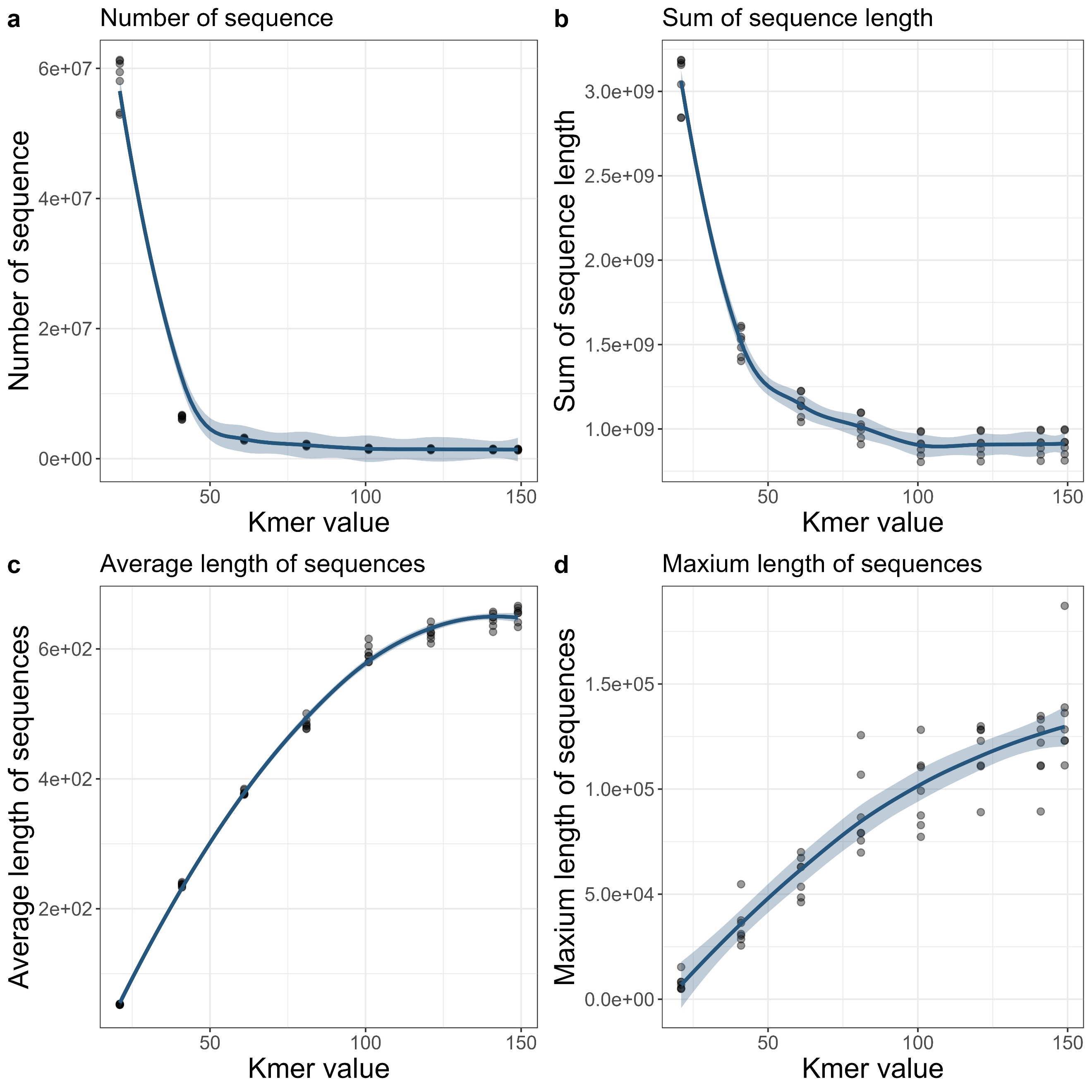

Figure S12. Statistics of intermediate assembly graph under different Kmer lengths (21, 41, 61, 81, 101, 121, 141, 149). The Kmer value = 41 was used for the intermediate assembly graph since more sequence information can be preserved and the moderate length of intermediate contig can be achieved. Seven samples were randomly selected for this analysis.

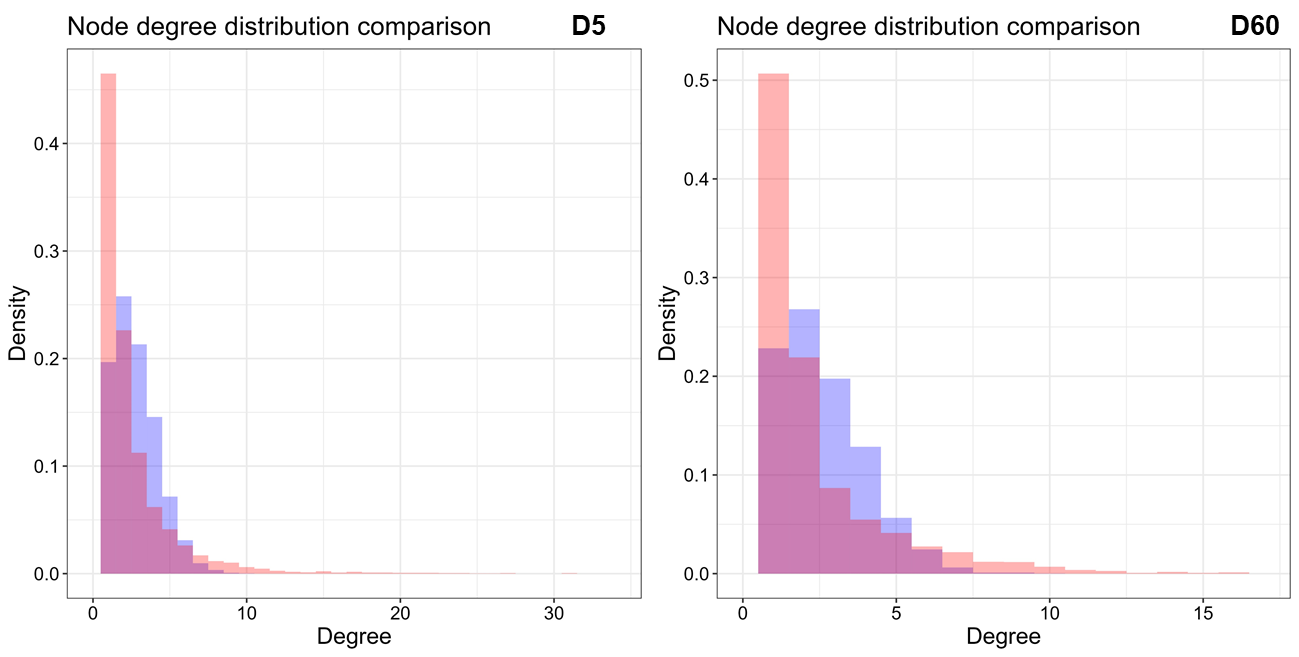

Figure S13. Comparison of node degree distribution between empirical (red) and random (blue) networks for D5 and D60.

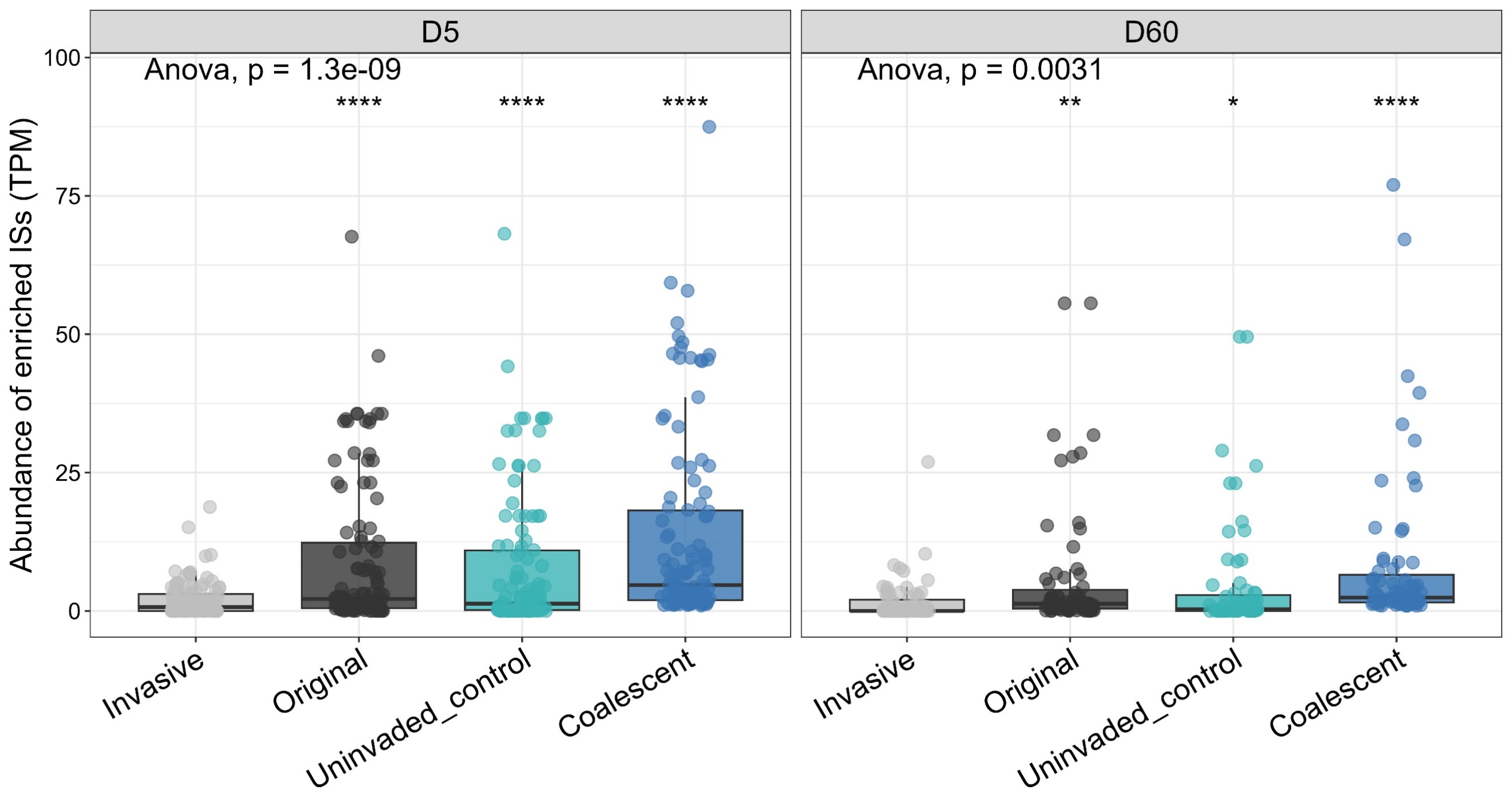

Figure S14. The abundance (TPM) of enriched ISs in each community. The difference in abundance between invasive and other communities was assessed by paired t-test and indicated with asterisks above boxes (“ns,” *p* > 0.05, “*,” *p* < 0.05, “**”, *p* < 0.01, “***,” *p* < 0.001, “****,” *p* < 0.0001). The lower relative abundance of enriched ISs in invasive communities suggested its weak contribution to the increase of ISs abundance in the corresponding coalescent communities when only 5% of invasive taxa were introduced for coalescences.

***Supplementary tables***

Table S1. The information of sequenced clean data.

| Sample Name | Clean Reads | Clean Base | Read Length | Q20 (%) | GC (%) |
| --- | --- | --- | --- | --- | --- |
| D0_1 | 39908513 | 11972553900 | PE150 | 95.03 | 60 |
| D0_2 | 33709194 | 10112758200 | PE150 | 95.1 | 60.44 |
| D0_3 | 40103609 | 12031082700 | PE150 | 95.44 | 60.56 |
| D0_4 | 40036947 | 12011084100 | PE150 | 95.82 | 59.04 |
| D0_5 | 40117905 | 12035371500 | PE150 | 96.19 | 59.69 |
| D0_6 | 40027733 | 12008319900 | PE150 | 95.66 | 59.39 |
| D0_13 | 40143708 | 12043112400 | PE150 | 96.15 | 57.69 |
| D0_14 | 40131461 | 12039438300 | PE150 | 96.36 | 57.14 |
| D0_15 | 40131342 | 12039402600 | PE150 | 96.33 | 57.56 |
| D0_22 | 40218147 | 12065444100 | PE150 | 95.51 | 59.5 |
| D0_23 | 40276352 | 12082905600 | PE150 | 95.68 | 59.3 |
| D0_24 | 40109215 | 12032764500 | PE150 | 95.79 | 60.21 |
| D0_25 | 40163957 | 12049187100 | PE150 | 95.87 | 59.71 |
| D0_26 | 40188088 | 12056426400 | PE150 | 95.79 | 60.26 |
| D0_27 | 40119166 | 12035749800 | PE150 | 96.03 | 61.2 |
| D0_28 | 40208848 | 12062654400 | PE150 | 95.95 | 61.07 |
| D0_29 | 40183982 | 12055194600 | PE150 | 95.84 | 60.27 |
| D0_30 | 40051148 | 12015344400 | PE150 | 95.65 | 60.29 |
| D5_1 | 40135659 | 12040697700 | PE150 | 94.99 | 60.67 |
| D5_2 | 40258047 | 12077414100 | PE150 | 95.26 | 60.33 |
| D5_3 | 40116035 | 12034810500 | PE150 | 95.59 | 60.75 |
| D5_4 | 40186203 | 12055860900 | PE150 | 95.33 | 60.15 |
| D5_5 | 40232107 | 12069632100 | PE150 | 95.35 | 60.38 |
| D5_6 | 40040184 | 12012055200 | PE150 | 95.23 | 60.41 |
| D5_13 | 40216257 | 12064877100 | PE150 | 95.15 | 60.21 |
| D5_14 | 40062647 | 12018794100 | PE150 | 95.38 | 60.39 |
| D5_15 | 36390426 | 10917127800 | PE150 | 95.28 | 60.27 |
| D5_22 | 28433815 | 8530144500 | PE150 | 95.01 | 60.27 |
| D5_23 | 38410393 | 11523117900 | PE150 | 95.09 | 60.43 |
| D5_24 | 40146556 | 12043966800 | PE150 | 94.99 | 60.29 |
| D5_25 | 40197352 | 12059205600 | PE150 | 95.69 | 60.61 |
| D5_26 | 40146764 | 12044029200 | PE150 | 95 | 60.25 |
| D5_27 | 39385444 | 11815633200 | PE150 | 95.14 | 60.31 |
| D5_28 | 39668398 | 11900519400 | PE150 | 95.13 | 60.52 |
| D5_29 | 40262639 | 12078791700 | PE150 | 95.1 | 60.33 |
| D5_30 | 40184836 | 12055450800 | PE150 | 95.31 | 60.57 |
| D60_1 | 40259420 | 12077826000 | PE150 | 95.23 | 59.37 |
| D60_2 | 40203599 | 12061079700 | PE150 | 95.48 | 60.38 |
| D60_3 | 40109445 | 12032833500 | PE150 | 95.27 | 60.77 |
| D60_4 | 40032362 | 12009708600 | PE150 | 95.11 | 60.73 |
| D60_5 | 40229826 | 12068947800 | PE150 | 95.78 | 60.56 |
| D60_6 | 40252213 | 12075663900 | PE150 | 95.47 | 60.63 |
| D60_13 | 40203060 | 12060918000 | PE150 | 95.11 | 60.27 |
| D60_14 | 40247925 | 12074377500 | PE150 | 95.8 | 60.54 |
| D60_15 | 40183788 | 12055136400 | PE150 | 95.08 | 60.47 |
| D60_22 | 40100801 | 12030240300 | PE150 | 95.07 | 60.33 |
| D60_23 | 40190895 | 12057268500 | PE150 | 94.92 | 60.11 |
| D60_24 | 38986439 | 11695931700 | PE150 | 95.2 | 59.21 |
| D60_25 | 37149653 | 11144895900 | PE150 | 95.07 | 59.72 |
| D60_26 | 40161014 | 12048304200 | PE150 | 95.28 | 60.11 |
| D60_27 | 40199570 | 12059871000 | PE150 | 95.23 | 59.87 |
| D60_28 | 40056178 | 12016853400 | PE150 | 95.07 | 60.18 |
| D60_29 | 40137695 | 12041308500 | PE150 | 95.27 | 61.4 |
| D60_30 | 40228012 | 12068403600 | PE150 | 95.12 | 60.34 |
